# Failure to classically condition planarian flatworms

**DOI:** 10.64898/2026.04.18.719399

**Authors:** Zachary S. Kelso, Madeleine C. Snyder, Samuel J. Gershman

## Abstract

Planarian flatworms represent one of the most evolutionarily informative nervous systems for an account of ancient bilaterian brains. Likewise, the unparalleled regenerative ability of planarians makes possible certain investigations of neural development, memory, and behavior that are simply impossible with other model organisms. Despite these facts, learning and memory are today underexplored in planarians, likely due in part to the controversial legacy of 20th-century experiments on the transfer of memories between individual flatworms. Here, we attempted to replicate and extend the classical conditioning experiments in planarians that were the basis of the later memory transfer work. We failed to find evidence of learning using both historical and contemporary conditioning protocols, and using computer vision methods to avoid subjectivity in manual video annotation. Our results cast doubt on the suitability of historic conditioning protocols for *Girardia dorotocephala* and *Schmidtea mediterranea* planarians. We encourage future researchers to explore more ethologically relevant conditioning paradigms to establish a behavioral foundation for investigating the molecular basis of learning.

## 1 Introduction

Evolutionary and comparative neuroscience suggest that planarian nervous systems represent the most basal, Urbilaterian-like brains of any contemporary organism [1–4]. Consequently, planarians offer a window into the evolutionary origins of cognitive brain functions, including learning, memory, and decision making. The twentieth-century discovery that planarians appear to be capable of learning and memory was a watershed in this research program. Even more remarkably, reports of memory retention after decapitation and regeneration, as well as memory transfer between individuals after RNA injections or cannibalism, suggest that planarian brains could shed light on novel mechanisms of memory storage. However, these claims have been mired in criticism and controversy, and contemporary research on planarian learning and memory is limited. The goal of this paper is to revisit basic questions about planarian learning in an effort to establish a behavioral foundation to further investigate these novel memory mechanisms.

Historically, planarian learning and memory has been tested with various forms of classical and operant conditioning [5, 6]. The seminal experiment was conducted by James McConnell and Robert Thompson in 1955 [7], after which McConnell became the leading force in planarian learning and memory transfer experiments [8–11]. These classical conditioning procedures were typically conducted with *Girardia dorotocephala* flatworms and employed light and shock as the conditioned (CS) and unconditioned (UCS) stimuli, respectively. All studies measured learning by quantifying turn and contraction responses to the onset of the CS after repeated stimulus pairings, and emphasized that the contraction response is more reliable than turns alone or pooled turns and contractions [5, 6, 12]. Allan Jacobson extended these experiments with additional controls and improved procedures [12–15], after which Jacobson remarked that “[i]t appears reasonable to assert that by the strictest criteria, classical conditioning has been conclusively demonstrated in planarians” [12]. Nonetheless, several groups that attempted replication failed to classically condition planarians or did so with only weak levels of learning. Though a review by Corning and Kelly [6] counted 36 inductions of classical conditioning using the original light-shock paradigm as measured by conditioned response acquisition, these were weighed against seven published failures, which included the mostly unsuccessful efforts of biochemist and Nobel laureate Melvin Calvin [16, 17]. Other forays into planarian learning also yielded negative results [18]. By the end of the 1970s, most planarian psychologists, including McConnell, had shifted their area of research [19]. The procedural ambiguities of how to reliably train and elicit conditioned behavior from planarians were never clarified, perhaps due in part to the reluctance of subsequent planarian researchers to associate themselves with the controversial legacy of the memory transfer episode [20].

Modern attempts to classically condition or otherwise train planarians have been few in the years since the initial movement [21–27], even as planarians became a fixture of regenerative biology [20, 28–32] and the subject of some neuroanatomical and pharmacological interest [33–36]. Research into planarians’ cognitive capacities has been renewed by recent investigations into non-synaptic forms of memory [37–42], which has returned attention to the McConnell-era transfer experiments and apparent ability of planarians to remember learned information even after decapitation and whole-brain regeneration [43, 44]. Nonetheless, the foundational classical conditioning experiments have not been exactly replicated. This is surprising given the volume of the historical literature, the utility of planarians as a model of evolutionarily ancient brains, and the disruptive implications of regenerative memory retention to the neuroscience of memory.

We therefore sought to replicate the original planarian conditioning experiments in alignment with Allan Jacobson’s protocol [13, 14]. We selected this particular protocol because it included the most rigorous set of controls (pseudoconditioning) to disambiguate true conditioning from sensitization or habituation (Methods). We refer to this protocol and our reproduction of it as the “historic” paradigm. We then tested a contemporary conditioning paradigm inspired by aversive conditioning in rodents (referred to as the “contemporary” paradigm), as well as a selection of ethologically-motivated alternative stimulus pairings (“variant” paradigms). In all cases, we were unable to find evidence for classical conditioning in either *Girardia dorotocephala* or *Schmidtea mediterranea* planarians. To address the possibility that these null results arose from the subjectivity of manual scoring (with high interscorer variability), we built a computer vision pipeline to analyze worm behavior across several days of training. The results of the computer vision analyses were likewise null.

## 2 Results and Discussion

In what follows, we describe our findings from attempting to classically condition *G. dorotocephala* flatworms using (1) a historic protocol based on the work of Allan Jacobson [12–15], (2) a contemporary protocol based on the work of James et al. (2026) [26], and (3) variant stimulus pairings informed by recent studies on planarian sensory capacities [13, 26, 45–47]. While our search space for a classical conditioning paradigm in planarians is not exhaustive, our assays show that salient stimuli across multiple sensory domains fail to elicit an increase in conditioned response rates comparable to those found in the original studies. We found no difference in conditioned response acquisition for any paradigm across two species: *Girardia dorotocephala*, which were historically used for psychological experiments, and *S. mediterranea*, which are commonly used in contemporary regenerative biology research. Finally, we found no difference in conditioning across three separate *G. dorotocephala* lineages from multiple locations across the United States (see Methods for more details).

### 2.1 Historic light-shock conditioning paradigm fails to produce conditioned response in planarians

We recreated the historic light-shock conditioning paradigm, in which *G. dorotocephala* planarians are subjected to two sessions of light habituation followed by five sessions of paired exposure to light and shock, remaining as faithful to the original protocol as possible with modern equipment (Fig. 1A-B; Methods). The classical conditioning group (CC; also called forward conditioning) received repeated pairs of light stimuli ending with an overlapping shock stimulus; the pseudoconditioning group (PC) received the same number of exposures to light and shock, but in a randomized and nonoverlapping order. This protocol controls for non-associative sensitization: if pseudoconditioned worms acquire a response to light comparable to that of the CC group, the response increase can be seen as an indication of sensitization rather than a stimulus-specific conditioned response or learned behavior.

**Fig 1.**
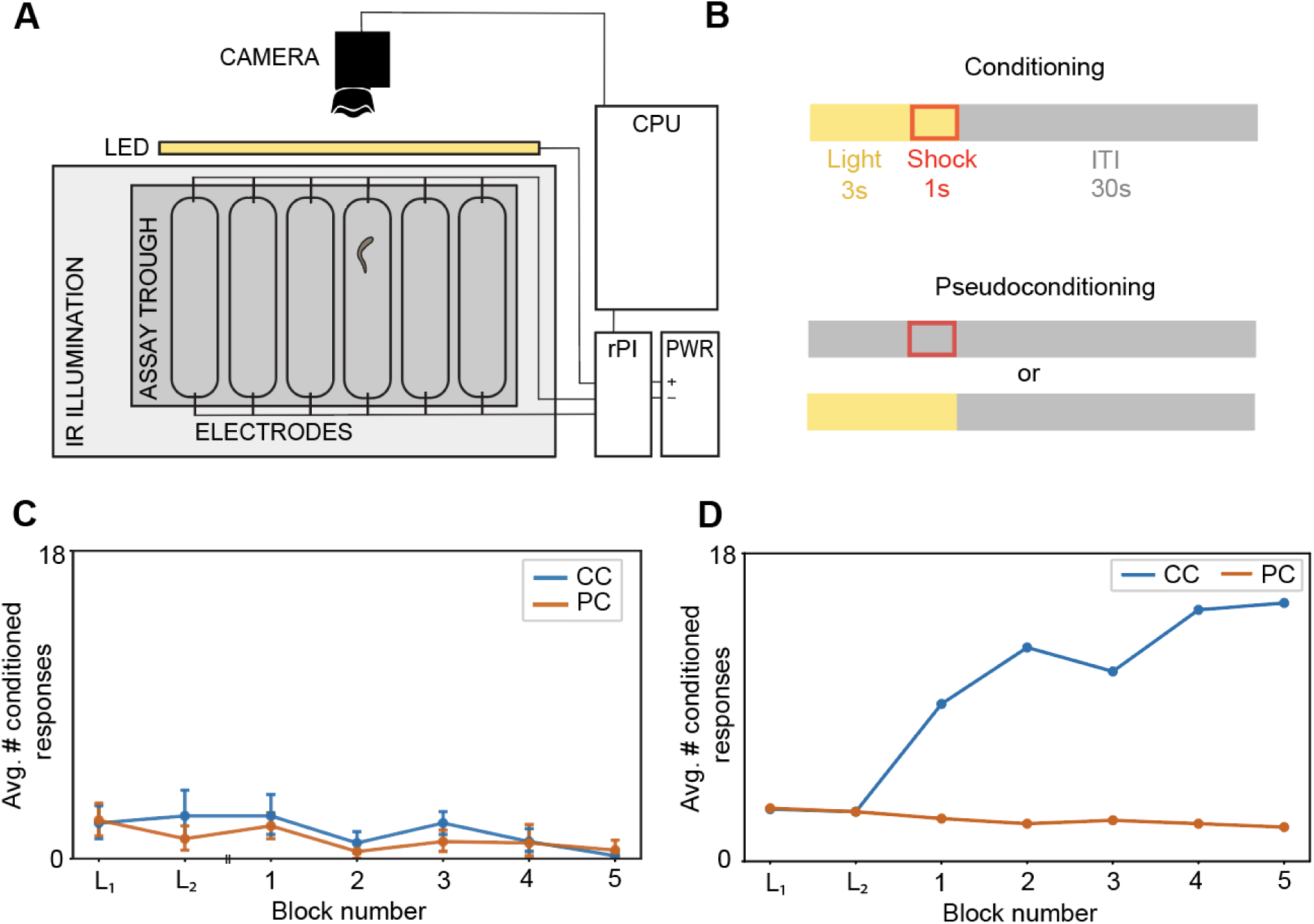
Recreation of historic conditioning paradigm. **A**. The experimental apparatus was a plastic trough of six parallel lanes with electrodes on the ends of each lane. A light source above the trough delivered the CS (white light), and an IR LED panel below the trough provided imaging illumination invisible to the planarians. A camera mounted above the trough recorded the experiments and a raspberry-Pi-based controller (rPI) controlled the delivery of CS and UCS (shock) stimuli in conjunction with a computer (CPU) and power supply (PWR). **B**. Each CC trial consisted of a three-second light CS that co-terminated with a one-second electric shock UCS (overlap shown by the red shock window within the yellow light window), followed by a 30-s intertrial ITI. In the PC group, the CS and UCS were delivered separately as unpaired shock-only trials or unpaired light-only trials, randomly interleaved (shown stacked). **C**. Mean number of conditioned responses across blocks, pooled across cohorts of *G. dorotocephala* worms. Blocks *L*_1_ and *L*_2_ are CS-only baseline blocks; blocks 1-5 are CC blocks. Blue line indicates CC cohorts (n = 12 worms); orange line indicates PC cohorts (n = 12 worms). Error bars are 95% CI. Both groups remained near baseline throughout conditioning, with no upward trajectory to suggest learning. The two groups did not differ from each other in overall response level. A stratified breakdown of the same data by baseline-responsiveness is presented in Figure S2A. **D**. Replotted from five published block means in Jacobson et al. (1967). The CC group shows a substantial, monotonically increasing response rate absent from our data. Original figure reprinted in Fig. S3.

In keeping with the historic experiments’ methods, observed planarians were hand-scored for contraction responses from the onset of the CS to the onset of the UCS (two-way interscorer percent agreement: 91.92%, but see Results 2.6). We found no significant increase in the number of conditioned responses per block of light stimuli across days; rather, we observed a significant decrease in the conditioned response rate across days (two-sided per-worm slope test: *t*(11) = −3.86*, p* = .003*, n* = 12 worms; Page’s *L* = 491.0*, p* = .998; Fig. 1C). There was no significant difference in conditioned responses between the CC and PC conditions (ANOVA main effect of condition: F(1, 22) = 2.88*, p* = .104). These results were markedly different from the stark increase in conditioned responses observed in the original study (Fig. 1D; original figure reprinted in Figure S3): the mean per-worm slope for our CC group was −0.02 ± 0.01 SEM responses/block, compared to Jacobson’s +1.40 ± 0.40 SE responses/block (Welch’s *t*(3.0) = −3.53*, p* = .039, propagating the SE of Jacobson’s 5-block-mean regression; 95% CI on the slope difference [−2.70, −0.14], excluding zero). The failure to replicate this result using the same species, stimulus parameters, apparatus, and pseudoconditioning controls suggests that either a meaningful phenotypic change in this species of worms has occurred since the original experiments, or that there is a critical but undocumented methodological detail that our recreation of the classic experimental setup fails to capture.

### 2.2 Individual differences in sensitivity to conditioned stimulus do not affect acquisition of the conditioned response

To investigate whether individual variability in response sensitivity to the conditioned stimulus may have obscured evidence of learning [48], we assayed six groups of six worms each with repeated presentations of the conditioned stimulus alone, then sorted them into low and high responders (Fig. S2A). The groups had a significantly different probability of reacting to the conditioned stimulus (white light) presentation (independent samples *t* -test, *t*(22) = 5.58*, p < .*001). We found no significant upward trend in either of the historically standard conditioned responses (turns or contractions) [5, 12] for both the conditioning group (which showed a significant decrease, as reported above) or pseudoconditioning group (*t*(11) = −1.82*, p* = .097), and slopes did not significantly differ between high and low responders in either the conditioning (Welch’s *t*(10) = −0.27*, p* = .794*, d* = −0.16) and the pseudoconditioning group (Welch’s *t*(10) = +0.51*, p* = .619*, d* = +0.30) (Fig. S2B). To minimize the risk of latent inhibition arising from stimulus pre-exposure during the responsiveness assay, we spaced the assay and the conditioning paradigm 2–3 days apart, during which the worms were only exposed to ambient light. Across the groups with high enough response numbers to discern from zero, we found no significant habituation to the conditioned stimulus (but see [6], Table VIII). The mean response rate increased significantly from Day 1 (11.8% ± 2.0% SEM) to Day 2 (17.5% ± 2.5% SEM; paired *t*(35) = 2.39*, p* = .022, two-sided; Cohen’s *d_z_* = 0.40). While this increase was consistent across five of six cohorts, no individual cohort reached significance alone. Slope analysis at the trial level did not reveal any significant gradual trend across trials (mean slope = 0*, t*(35) = 0.16*, p* = .873), suggesting the increase reflected a between-session shift rather than a continuous within-session change (Fig. S2C). These findings rule out individual variation in baseline CS sensitivity as an explanation for our overall null result.

### 2.3 Feature-based quantification of the conditioned response in contemporary paradigm fails to provide evidence for learning

After the historic conditioning protocol yielded no appreciable evidence of learning, we attempted to replicate a contemporary conditioning protocol adapted from fear conditioning experiments in rodents [26] using *G. dorotocephala* and the distantly related planarian *S. mediterranea*, a common model organism in regenerative biology today [31, 32]. To quantify the conditioned responses in this protocol, we used both manual annotation of turns and contractions and a novel computer vision pipeline (Fig. 2A). From the hand-scored data we found that neither conditioned turns nor contractions showed a significant upward trend over training days in the conditioning group (turns: per-worm slope test, *t*(41) = −1.95*, p* = .97, one-sided; Page’s *L* = 1026.5*, p* = .90; contractions: *t*(41) = −0.96*, p* = .83; Page’s *L* = 1035.5*, p* = .78; *n* = 42 worms) or the pseudoconditioning group (turns: *t*(11) = −1.11*, p* = .86; contractions: *t*(11) = 0.35*, p* = .37; *n* = 12 worms). Linear mixed models (LMMs) confirmed no significant Day × Condition interaction for either turns (*β* = −0.18*, z* = −0.21*, p* = .83) or contractions (*β* = 0.12*, z* = 0.36*, p* = .72), indicating that the conditioning group did not produce responses at a different rate than controls. The two groups did differ in overall turn frequency (LMM: *β* = −7.65*, z* = −3.12*, p* = .002), with the conditioning group responding more than controls, but this difference was present from the outset and did not change with training (Fig. 2B). Breaking the conditioning group down by species and strain (Fig. S1) confirmed this pattern in every measurable sub-population: per-worm conditioned response slopes were significantly lower than the historic learning rate of +1.40 responses/day (Welch’s *t* -test, with denominator df adjusted to propagate the SE of Jacobson’s regression). By subpopulation and conditioning paradigm: *S. mediterranea* CC (−1.75 responses/day, *n* = 6, *t*(7.5) = −3.95*, p* = .005), *G. dorotocephala* (Michigan, USA) CC (−0.33, *n* = 6, *t*(7.5) = −3.06*, p* = .017), *G. dorotocephala* (stock) CC (−0.86, *n* = 30, *t*(21.3) = −3.08*, p* = .006), and the matched *G. dorotocephala* pseudoconditioning reference (−0.43, *n* = 12, *t*(11.2) = −2.95*, p* = .013); no sub-population produced a positive group-mean slope, and all were significantly lower than the Jacobson reference slope (Welch’s *t* -tests, all *p* ≤ .017).

**Fig 2.**
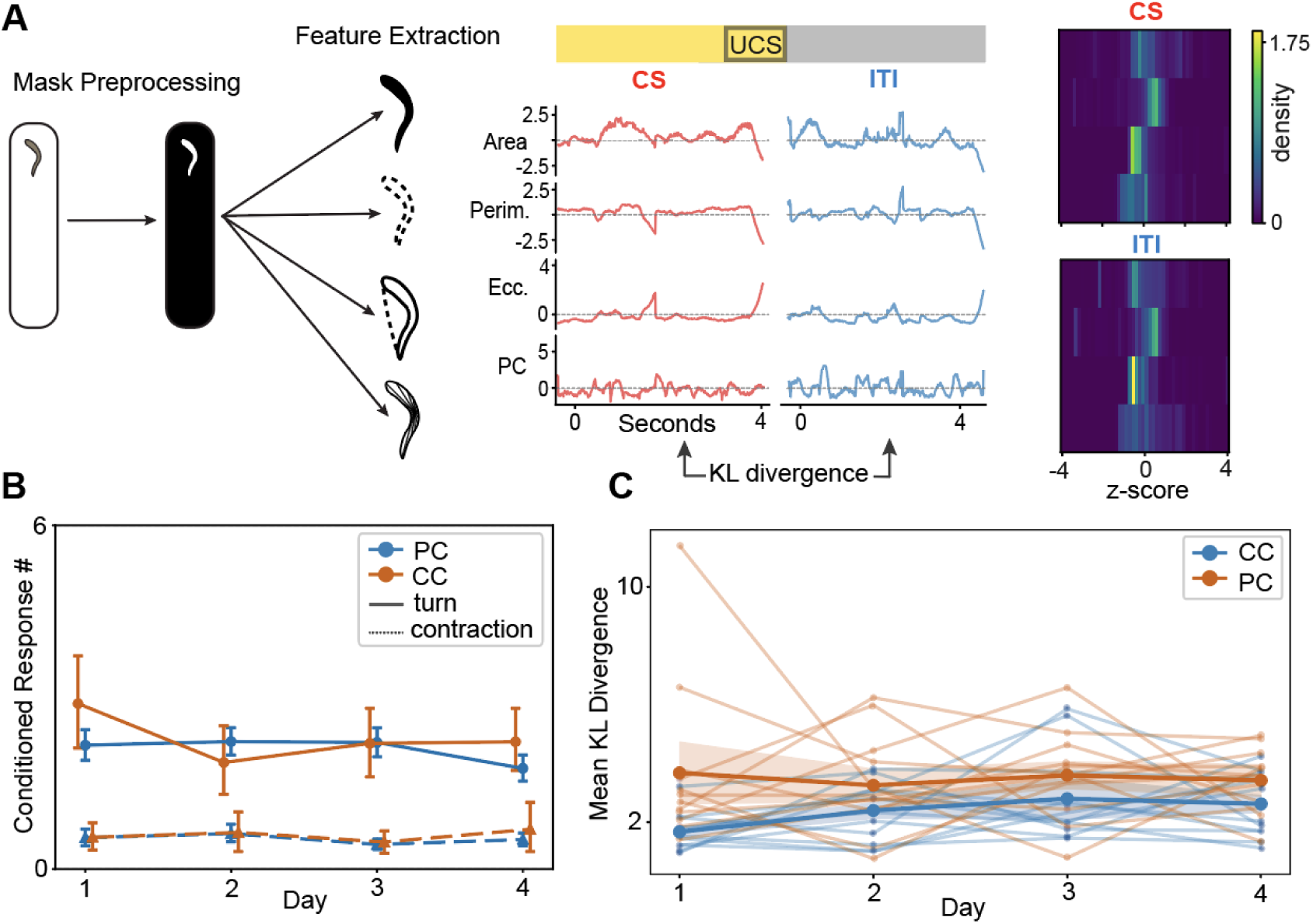
Application of machine vision pipeline to assess learning over the course of a contemporary conditioning protocol. **A**. Feature extraction pipeline. Four of ten morphological/spatial features shown (Area, Perimeter, Circularity, PC1). Traces show z-scored feature values for an example worm during CS-on (red) and ITI (blue) periods; heatmaps show corresponding feature distributions. Increasing area with stable perimeter indicates lateral body spread; spikes in circularity with decreasing perimeter indicate contraction. **B**. Hand-scored conditioned responses across training days (CC: *n* = 42, blue; PC: *n* = 12, orange), pooled across species and strains. Solid lines: turns; dashed lines: contractions. Error bars indicate 95% bootstrap CIs. Species-level breakdown in Fig. S1B. **C**. Computer-vision-derived KL divergence (CS-on vs. ITI) across training days for stock *G. dorotocephala* (CC: *n* = 12, blue; PC: *n* = 12, orange). Thin lines: individual worms; bold lines: group means ± 95% bootstrap CI. Note that neither group shows a systematic upward drift and PC trajectories are if anything slightly higher than CC trajectories at every day, in the opposite direction to what learning would predict, and the two group-mean trajectories overlap within their CIs at all four time points.

Because variability in subjective manual scoring could obscure genuine group differences, we built a custom feature-extraction pipeline using Segment-Anything (Meta Inc.) and shape-analysis methods previously applied to planarian behavior [45, 49]. Following a recently developed approach for comparing the acquisition of learned behavior across species and paradigms [50], we quantified worm behavior by calculating the Kullback–Leibler (KL) divergence between distributions of body-shape features sampled during conditioned-stimulus-on (CS-on) and intertrial (CS-off) intervals (Fig. 2A). Intuitively, the KL divergence measures how much the presence of the CS changes the distribution of morphological features such as body area or perimeter. A worm that consistently responds to light will look systematically different during CS-on periods than during the ITI, producing high KL divergence, whereas a worm that ignores the CS will look similar in both windows, producing low KL divergence. Consequently, if animals acquire a conditioned response over training, the divergence between CS-on and CS-off feature distributions should increase across days, and should do so more steeply in the conditioning group than in the pseudoconditioning group.

We fit a linear mixed model predicting log-transformed KL divergence from Day (continuous), Condition (conditioning vs. pseudoconditioning), and their interaction, with random intercepts for individual subjects. The interaction term was not significant (*β* = −0.08*, z* = −0.95*, p* = .342), indicating no evidence that the conditioning group’s trajectory differed from that of the pseudoconditioning group. This was corroborated by a direct comparison of per-subject slopes between groups (conditioning: +0.33 ± 0.13; pseudoconditioning: −0.04 ± 0.31; Welch’s *t* = 1.11*, p* = .286). The two groups did differ significantly in overall KL divergence (*β* = 0.35*, z* = 3.61*, p < .*001), though notably the pseudoconditioning group showed a higher mean KL divergence, which is opposite to what a conditioning effect would predict (Fig. 2C). The higher mean KL divergence in the pseudoconditioning group may reflect greater behavioral variability induced by unpredictable stimulus timing, though we do not have a principled account of this difference. Critically, the convergence of two independent measurement methods (manual scoring and automated computer vision) on the same null result substantially reduces the likelihood that the negative outcome reflects a measurement artifact.

### 2.4 Ethologically relevant stimulus pairings fail to evoke a robust conditioned response

Modern literature on planarian sensory capacity motivated us to substitute light and shock for ultraviolet (UV) light, which is a more ethologically relevant stimulus [45–47, 51]. Planarians have UV-sensitive extraocular opsins distributed across their bodies that mediate a rapid contraction in response to high-amplitude UV light exposure [46, 52]. Low-amplitude and slowly modulating UV light evoke a more subtle and complex response [36], and are suitable as conditioned stimuli. We selected two new stimulus pairings (CS and UCS): (1) white light and high-amplitude ultraviolet (UV) light, and (2) low-amplitude UV light and shock. We suspected that planarians may display implicit biases for detecting and associating these sets of stimuli, making it easier to elicit a robust conditioned response.

We first used low-amplitude UV light as the CS and shock as the UCS, reasoning that low-amplitude UV would evoke a salient yet suppressible response. In the historically used species (*G. dorotocephala*, n = 18 worms), the per-worm acquisition slopes across five conditioning blocks did not increase (mean slope = −0.07 ± 0.07 SEM responses/block, 95% CI [−0.22, +0.08]) (Fig. 3A). We then used white light as the CS and high-amplitude UV as the UCS, choosing the latter because planarians do not habituate to phasic, high-amplitude UV exposure [45]. Neither species we tested showed an upward trend (Fig. 3B); the per-worm slopes were −0.67 ± 0.44 SEM responses/block in *G. dorotocephala* (*n* = 6, 95% CI [−1.80, +0.47]) and −0.67 ± 0.49 SEM responses/block in *S. mediterranea* (n = 6, 95% CI [−1.94, +0.60]). Across all three protocol × species combinations, the per-worm acquisition slope was significantly lower than the slope reported by Jacobson et al. (1967) (Fig. 3C). The Jacobson CC reference slope, recomputed from his five published block means, is +1.40 ± 0.40 SE responses/block. Two-sided Welch’s *t* -tests comparing each of our populations to the Jacobson reference rejected equality at *α* = 0.05 in every case: GD low-UV/shock *t*(3.2) = -3.59, p = .034 (95% CI on the difference [−2.72, −0.21]); GD white/high-UV *t*(7.8) = −3.46, *p* = .009 ([−3.45, −0.68]); SM white/high-UV *t*(8.0) = −3.24, *p* = .012 ([−3.54, −0.60]). In each of our populations the 95% CI excluded Jacobson’s point estimate of +1.40, and the standardized effect against the fixed Jacobson reference was large (one-sample Cohen’s *d* = -4.91, -1.91, and -1.71, respectively). These results indicate that neither variant using UV light replicated the conditioning rate originally reported by Jacobson et al.

**Fig 3.**
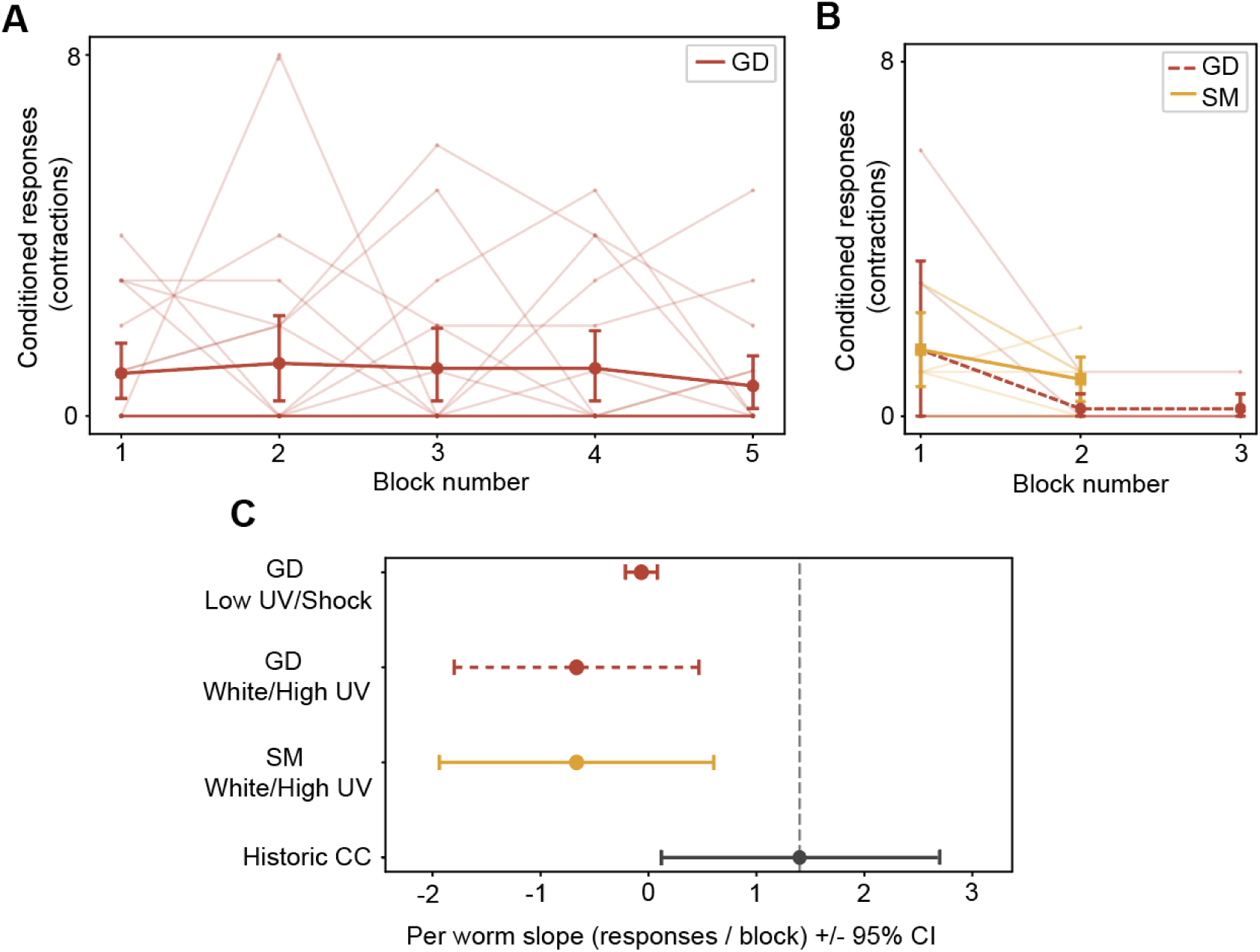
Failure to elicit learning using ethologically-motivated conditioning stimulus variants. **A**. Low-UV + Shock paradigm (*G. dorotocephala*, *n* = 18). Thin red traces: individual worms; thick dark-red line: pooled mean ± 95% bootstrap CI. **B**. White + High-UV paradigm in stock *G. dorotocephala* (dashed red, *n* = 6) and *S. mediterranea* (solid yellow, *n* = 6), each completing the maximum number of conditioning blocks achievable for that species. Thin traces: individual worms; thick lines: group means ± 95% bootstrap CIs. **C**. Forest plot of per-worm OLS acquisition slopes (filled circles, 95% CI) versus the historic reference (+1.4 responses/block, dashed vertical line). Historic CC slope recomputed by OLS on five published block means (df = 3). Note that all groups’ slopes are significantly lower than the historic learning rate.

### 2.5 Planarian species and origin have no observable effect on learning

Literature from the 1960s claimed that worm origin, species, and feeding interval (among many other factors) are important for the success of planarian learning [53]. To test whether worm provenance and species affected our worms’ learning outcomes, we collected *G. dorotocephala* specimens from three locations across the United States, including the location (to the best of our knowledge) where McConnell procured his population of planarians for most (but not all) of his experiments. Commercially available *G. dorotocephala* from North Carolina (also called stock planarians), wild-caught Michigan *G. dorotocephala*, wild-caught California *G. dorotocephala* and laboratory *S. mediterranea* worms were used across our previously described experiments and showed indistinguishable levels of responding. While different species demonstrated notably different levels of general activity, we did not find that a particular species had an effect on the acquisition of conditioned responses (Fig. 3A; Fig. S1). Laboratory *S. mediterranea* are from the Mediterranean [54], whereas *G. dorotocephala* are native to North America, and their evolutionary distance is possibly as vast as 100 million years [55]. Thus, the failure of American (Michigan, California, and Carolina strains) and Mediterranean species to demonstrate learning of conditioned responses is more consistent with a paradigm-level failure than with any species-specific deficit.

### 2.6 Behavioral scores are not consistent across trained observers

Having found no evidence of learning as measured through increases in conditioned response rates, we endeavored to determine the extent to which different scorers could agree on what types of behavior count as turns and contractions. For the historic paradigm, trials were scored in a binary fashion and 91.92% of trials were given the same value (“0” for no contraction or “1” for a contraction) by both scorers. This agreement value is dominated by trials where each scorer agreed that no response had occurred (89.67% for “0-0” and only 2.25% for “1-1”). When looking only at trials where at least one scorer reported a positive response, the interscorer agreement is 21.78%. For all trials, including those rated “0-0”, Cohen’s *κ* = 0.31*, P*_0_ = 0.92, and *P_e_* = 0.88. A reliable, robust assessment of contractions does not follow from these data.

In the contemporary paradigm, rather than a binary assessment of contraction, behavior was scored as counts of turns and contractions across the active duration of the conditioned stimulus, so a more complex metric of interscorer agreement was required. Three scorers scored a sample of 203 videos of our footage and timestamps of turns and contraction events were analyzed (see Methods). While the total sums of turns and contractions for each video are essentially the same for each scorer (Fig. S4), an analysis of the actual timestamps of reported responses reveals severe disagreement. The primary result is that unanimous agreement of contraction events (all scorers marked a given event in a given video as a contraction) is only 28.15% (67/238 of all contraction events). The agreement for turn events is even lower at 15.14% (106/700 of all turn events). These results echo those of Arlene Hartry [18], who similarly examined unanimous contraction agreement between three observers of planarian conditioning footage and found that so “few responses fell into this category” that they were statistically irrelevant. The low agreement of multiple scorers is in marked contrast to conventional classical conditioning procedures, where it is understood that conditioned responses should be relatively obvious to the trained eye.

To further examine interscorer agreement at the timestamp level, we computed F1 metrics for each pairwise combination of the three scorers’ annotations for each video. The F1 metric is the harmonic mean of precision and recall measures (Methods). In the context of our data, precision is the number of the Test Scorer’s events (turns or contractions) that the Reference Scorer also marked. Recall is the proportion of Reference events that the Test Scorer correctly identified out of all of the Reference’s events. These proportions describe how much the Test Scorer is able to capture a Reference Scorer’s data while minimizing the detection of non-reference events. If a pair of scorers agreed perfectly, an F1 score of 1 would be expected, and severe disagreement would yield a 0. We also quantified event confusion based on the proportion of unpaired Reference events that have a temporally proximate Test event of the opposite type in the test events . These instances occur when the Test and Reference Scorer recognize the same event as having occurred, but categorize it differently.

Table 1 shows the recall, precision, F1, and confusion rates for each scorer pair and event type. Broadly, these measures indicate poor agreement about when events occur across scorers: on average, a given scorer detects about half of the turns (52.3%) or contractions (59.7%) that another scorer marks, and produces an equal amount that their reference fails to detect (55.6% for turns, 57.8% for contractions). To compare contraction agreement in response to the conditioned stimulus (light) versus the unconditioned stimulus (electric shock), Scorer 1 and Scorer 2 additionally examined 198 clips from the same set of videos for responses during the shock period. For shock-induced contractions, Scorer 1 and Scorer 2 agreed on 86.87% of trials (172/198). The discrepancy between high agreement about shock-induced contractions but low agreement about light-induced contractions fails to provide evidence that the CS-period contractions are specific, learned responses to light and not just ambient, noisy movement. This interpretation is supported by the observation that the total number of observed turns and contractions is essentially consistent across scorers despite starkly different timestamp attributions for those behaviors (Fig. S4). The sum of these results suggests that manual assessment of planarian behavior is not robust for our experimental footage. That is, when multiple scorers observe and describe the same experimental footage, their reports are not reliably similar, even if the raw counts of turns and contractions are stable across scorers.

**Table 1.** Pairwise comparison of scorer agreement across behaviors as measured by recall, precision, and F1 scores of event timestamps of 203 sample training videos. Note that when a Reference and Test Scorer are flipped, the recall and precision values likewise swap places, but the F1 score remains the same.

| Behavior | Reference | Test | Recall | Precision | F1 | Confusion Rate |
| --- | --- | --- | --- | --- | --- | --- |
| Turns | Scorer 1 | Scorer 2 | 0.540 | 0.745 | 0.626 | 0.029 |
| Turns | Scorer 2 | Scorer 3 | 0.397 | 0.640 | 0.490 | 0.009 |
| Turns | Scorer 3 | Scorer 1 | 0.632 | 0.284 | 0.392 | 0.000 |
| **Turns** | **Average** | – | **0.523** | **0.556** | **0.503** | **0.013** |
| Contractions | Scorer 1 | Scorer 2 | 0.827 | 0.602 | 0.697 | 0.000 |
| Contractions | Scorer 2 | Scorer 3 | 0.492 | 0.556 | 0.522 | 0.031 |
| Contractions | Scorer 3 | Scorer 1 | 0.473 | 0.576 | 0.519 | 0.178 |
| **Contractions** | **Average** | – | **0.597** | **0.578** | **0.579** | **0.070** |

Our finding of low interscorer agreement is concerning for the interpretation of the historic literature on planarian conditioning, all of which relies on manual scoring. Though several of the classic studies do report high interscorer agreement, in many cases the complete data, the precise interscorer methods, and the experimental footage have been lost or are simply not available for review. The inconsistency in manual scoring of conditioned responses suggests, at least for our data, that we are attempting to quantify an absent learning signal, which is a poor foundation upon which to investigate potentially paradigm-shifting questions about memory transfer and the molecular basis of memory.

### 2.7 General Discussion

Across three conditioning protocols, two species, four geographic populations, and two independent measurement methods, we found no evidence for the robust classical conditioning of freshwater planarians reported in historic literature. The consistency of this null result makes a simple procedural explanation unlikely. We suggest that the conditions for eliciting robust and reproducible conditioned behavior in the tested planarian species are highly idiosyncratic, and we encourage future research to explore more ethologically guided stimuli that are more likely to produce robust learning behaviors. Because of our failure to demonstrate conditioned behavior, we were not able to address questions of molecular transfer of memory between planarians in any way.

It is difficult to reconcile our null results with the 20th century planarian learning literature. Several laboratories (not only the laboratories of McConnell and his students) reported success in training planarians, and over fifty scientific papers were published reporting successful conditioning of planarians between 1959 and 1971 [6], including in high-impact journals like *Nature*. Additionally, editions of the *Worm Runner’s Digest* contain letters from students and laypeople describing their efforts to condition planarians with an approximately 4:1 ratio of success to failure [17]. Publication bias toward positive results may partly explain the apparent preponderance of successes in the historic literature, though this cannot be the whole story given that different labs independently replicated a positive result. Other demonstrations of apparent learning include instrumental and maze paradigms (e.g. [56–58]), which we have not attempted to replicate here. Modern literature suggests that *C. elegans*, which have far fewer neurons than planarians (302 in hermaphrodites compared to 10,000-12,000 in the planarian brain) are capable of aversive learning [59–62]. We believe our efforts do not reflect on the widespread ability of planarians to learn in any situation, but conclude that we were unable to specifically elicit classical conditioning in the historically championed species *G. dorotocephala* or commonly used species *S. mediterranea* using historic and contemporary conditioning paradigms. Ultimately, we cannot explain why our results are in such contrast to those in the historic literature.

We close with a quote from a 1964 paper by Bennett and Calvin [16], who attempted to train planarians on a variety of tasks and were mostly unsuccessful:

> “Until more adequate and more useful methods and descriptions of methods to train planarians are available, this animal appears to have little utility for proposed studies of the possible biochemical basis of memory.”

This assessment appears, at least in our hands, to hold true today. Going forward, we suggest that researchers methodically document their procedures, data, and footage demonstrating learned behavior in the context of the selected conditioning protocol. Perhaps more importantly, we recommend exploring alternative stimulus combinations informed by planarian ethology and ecology. Despite the failure to replicate historic results, we are optimistic that future work will discover a robust behavioral foundation for conditioning, providing an unprecedented model through which to study neural circuit development and the physical basis of learning and memory.

## Supporting information

Supplementary information - revised

## Acknowledgments

We extend our gratitude to the laboratories of Peter Reddien and Michael Levin for supplying worms and providing technical guidance, and to the Harvard NeuroTechnology Core for aid in building apparatuses. We additionally thank Reeva and Daniel Kimble for discussions of their experiences training planarians in the laboratory of James McConnell in the 1950s and 1960s, and for generously providing us with access to the complete *Worm Runner’s Digest*.

## Data and Code Availability

Code to reproduce the main and supplementary figures in this manuscript is available here: https://github.com/madeleinesnyder/PlanariaPipe/tree/main and the mp4 files used for manual scoring and binary masks used for the feature extraction pipeline are available on the Harvard Dataverse (https://doi.org/10.7910/DVN/NSTJGZ). The .bin files used to create the masks using our extraction pipeline are available upon request due to storage limitations.

## 3 Materials and methods

There are three major paradigms in this study. The first is our attempt to reproduce the classic 20th century planarian light-shock conditioning experiments using a similar apparatus and set of experimental parameters as those used by James McConnell and Allan Jacobson (stimulus timing, planarian species, etc.). We refer to this as the “historic” classical conditioning paradigm. The second paradigm we call “variant” because they represent an exploration of novel stimulus configurations and timing parameters that are unprecedented in the planarian memory literature. Third comes the “contemporary” paradigm, which seeks to replicate a recent paper that used a spaced conditioning regime adapted from rodent experiments to classically condition planarians. The particular details of these three paradigms are summarized in Table 2 and given in detail below.

**Table 2.** Different paradigms used in this study. **Table 2**. Tabulation of each classical conditioning paradigm’s experimental parameters for our attempted replications of the historic protocol, our variations of the historic protocol with alternative stimuli, and the contemporary protocol.

|  | Historic replication | Variant paradigms | Contemporary replication |
| --- | --- | --- | --- |
| **Role in study** | Attempted replication of historic Jacobson [13,14] studies | Explorations of alternative stimuli that might more readily elicit learning | Attempted replication of recent planarian conditioning work based on rodent conditioning [26] |
| **Species** | *G. dorotocephala*, Carolina strain | *G. dorotocephala* of multiple strains and *S. mediterranea* | *G. dorotocephala* of multiple strains and *S. mediterranea* |
| **Apparatus type** | Parallel lanes with electrodes (PLA material) | Parallel lanes with electrodes (PLA material) | Parallel lanes with electrodes (milled plastic) |
| **CS** | 3 sec white light | White or ultraviolet light | 30 sec blue light |
| **UCS** | 1 sec AC shock, coterminating with CS | Coterminous 1 sec UV light or AC shock | 3 sec AC shock, coterminating with CS |
| **Control** | Pseudoconditioning by alternating light and shock | None | Pseudoconditioning by alternating light and shock |
| **Trial timing** | 125 trials in 5 blocks of 25 trials across 2 days | 25 trials per block in 3–5 blocks over 2–3 days | 24 trials in 8 blocks of 3 trials across 4 days (2 blocks/day) |
| **Intertrial interval** | 30 seconds | 2 minutes | 30 seconds |
| **Interblock interval** | 10 minutes | 10 minutes | 4 hours |
| **Scoring method** | Hand-scored | Hand-scored | Hand-scored and computer vision pipeline |
| **Other** | Before training, worms received 2 blocks of 25 light-only trials. | Control conditions not run due to low response in non-control worms | See Methods for procedural divergence from original study |

### 3.1 Planarian acquisition and care

Planarians were housed in 6- and 11-cup rectangular Pyrex dishes with air holes carved into the plastic lids. The dishes were filled to a level of 2–3 inches with Montjuic water (1.6 mM NaCl, 1.0 mM CaCl_2_, 1.0 mM MgSO_4_, 0.1 mM MgCl_2_, 0.1 mM KCl, 1.2 mM NaHCO_3_ in deionized water). Housing dishes were cleaned biweekly on Tuesdays and Fridays. Cleaning consisted of corralling planarians into one corner using a water jet created by a transfer pipette or spray bottle containing fresh Montjuic water, draining the water from the dish, and wiping down the floor and walls with unbleached paper towels. The corner where the planarians had been concentrated was not wiped down, leaving an untouched triangular region with an area of 2–3 square inches. The unwiped corner was alternated between opposing sides with each cleaning so that a continuous zone of slime was always present in the trough. For experiments, planarians were individually isolated into six-well plates (Falcon, item no. 353046) containing approximately 10 mL of Montjuic water four days prior to the start of training.

Planarians were fed with freshly hatched *Artemia* brine shrimp nauplii (Brine Shrimp Direct, item no. BSEA8Z), hatched in salt water according to vendor instructions. For feeding, the hatchlings were rinsed with Montjuic water and transferred to the planarian dishes with a transfer pipette. Confirmation that the planarians ate the shrimp came in the form of pursuit of the shrimp, visible pharyngeal contractions, and the tendency for planarians to take on the color of their food (orange-red with brine shrimp). Planarians were fed once or twice per week, and in the case of experiments, starved for seven days prior to testing.

The planarians used in these studies varied in source. The historic experiments used worms purchased from Carolina Biological Supply (item no. 132954), which are wild-caught from various water sources in North Carolina. Miscellaneous experiments using alternative stimuli used the Carolina *G. dorotocephala* as well as *S. mediterranea* worms originating from the laboratory of Peter Reddien. Experiments modeled after the contemporary James et al. study used the Carolina *G. dorotocephala* worms and *S. mediterranea*, as well as a strain of wild-caught *G. dorotocephala* descended from an individual acquired from the Huron River in Ann Arbor, Michigan, where James McConnell is suspected to have sourced worms [63]. This founder planarian was found June 12, 2025, on the bottom of a rock in the shallow pool at the base of the Argo Dam on the outflow side (coordinates: 42°17’24.7”N, 83°44’42.6”W). Its species was determined visually by its prominent auricles and brown, unspotted coloration, and distinguished from the similar species *G. tigrina* by its comparatively weaker adhesion to glass. Once in the laboratory, this individual worm was amplified into a colony by repeated cycles of bisection and regeneration. In a similar fashion, two worms of the same species were acquired from a stream on the University of California, Berkeley campus, which were used in a variant (White-UV) experiment.

### 3.2 Assay chamber, stimuli, and data acquisition

We designed an assay chamber resembling side-by-side duplicates of McConnell’s original chamber [64]. Six separate roundbottomed, ovoid lanes were milled from a transparent block of polycarbonate and header pins were inserted at the ends of each lane. Sets of ipsilateral pins were connected by a copper wire to transmit the electrical shock stimulus to all lanes simultaneously (Fig. 1A). Two alligator clips attached to the pin-spanning copper wires connected each side of the trough to a DC power supply via an H-bridge module that together produced a 10-12 V, 60 Hz square-wave AC shock stimulus. While there is debate regarding the extent to which the orientation of a DC current affects conditioning in planarians [65], the primary historical study we are replicating used alternating current [14]. Experiments were recorded at 10 Hz with an overhead camera (Teledyne FLIR GS3-U3-23S6C-C (2048px x 2048 px) or GS3-U3-41C6M-C (4096px x 3000px)) suspended approximately one foot above the trough. A long-pass filter (750 nm) was placed over the camera lens to eliminate saturation effects from the light stimulus. The trough was placed on a white acrylic panel which diffused another panel of infrared (IR) LED lights (850nm, Waveform Lighting flexstrips) below, so the worms were clearly visible from the overhead camera. For the white light in the historical reproduction, we employed a 100-Watt incandescent bulb (Fluker’s Incandescent Basking Spotlight Bulb). Where ultraviolet light was a stimulus, a 100-Watt LED UV panel (395nm Glostars 100W LED, IP66) was used. In the case of the low ultraviolet condition, a diffuser was placed over the panel to reduce the light intensity. For the contemporary paradigm, we used an acrylic panel covered in rows of LED strips (6500K) with the panel itself covered in blue cellophane per the James et al. study. Small IR LEDs were directly coupled to the shock and light stimuli and visible in the camera frame to serve as indicators of stimulus onset for the viewer.

Custom software written in Python controlled the camera, conditioned stimulus, and unconditioned stimulus via a Raspberry Pi Pico (RP2040) microcontroller. Data were collected onto a Linux computer (Ubuntu 20.04.6 LTS, 64 bit) using a custom Python (version 3.8.10) recording script. The assay rig was contained in a rectangular box made of acrylic panels affixed to metal frame to isolate the experimental equipment and reduce ambient illumination.

### 3.3 Historic classical conditioning paradigm

In Allan Jacobson and colleagues’ experiments [13–15], planarians were rapidly and repeatedly presented with overlapping light and shock that resulted in the worms contracting to light even in the absence of shock. We attempted first to reproduce these experiments in their timing stimulus parameters. Planarians of the species *G. dorotocephala* (known as *Dugesia dorotocephala* in Jacobson’s time) were purchased from Carolina Biological Supply (item no. 132954) and subjected to 125 trials of 3 seconds of light, the final second of which included a 10-12 V, 60 Hz square-wave AC shock. Trials occurred in five blocks of 25 trials, each trial separated by 30 seconds, with ten-minute rest periods between blocks that were themselves spread over two days. Day 1 contained two consecutive light-only blocks in accordance with the historic paradigm. These were identical to the light-shock blocks, save that the shock did not occur. Presentation of the CS alone without any paired UCS had been reported to diminish the effectiveness of conditioning as far back as 1959 [66], but because the Jacobson group gave these light-only trials, we did likewise. The two Day 1 light-only blocks were followed by two light-shock blocks. On Day 2, worms received three light-shock blocks. In the case of the pseudoconditioning control, light-shock blocks were replaced with light-or-shock blocks. These consisted of randomly alternating light or shock presentation, with the constraint that no more than four consecutive light stimuli and no more than two consecutive shock stimuli were permitted.

Pseudoconditioning is nowadays often implemented by independent Poisson processes for different stimuli, but random alternation is the method for Jacobson’s own pseudoconditioning control, so as with the pretraining light-only blocks, we did likewise. The pseudoconditioning was implemented by generating random lists of 25 alternating L and S symbols, then filtering for those that violated the consecutivity restraints. A random candidate LS sequence was selected at the start of each pseudoconditioning block and each trial in that block corresponded to a symbol in the LS sequence. For example, a pseudoconditioning block that selected the sequence “LLS…” would present two light trials and then a shock trial. The training footage was scored on a trial-by-trial basis and conditioned responses were marked when a worm performed a “discernible longitudinal contraction of the body” [13] in the two-second period between light onset and shock onset. This paradigm, along with the others described below, is summarized in Table 2.

We diverged from Jacobson’s experiments in select ways consistent with established guidelines for training worms. First, before experiments began on any given day, a separate cohort of approximately 20 non-experimental worms were allowed to “slime” the trough. This was done in accordance with previous research that found that the presence of slime improved memory performance [53, 67]. After twenty minutes, the non-experimental slime worms were removed and experimental worms were allowed to acclimate in the trough for ten minutes before experiments began. After a cohort of worms was done for the day, the trough was rinsed with deionized water and scrubbed with unbleached paper towels. Second, whereas the historic Jacobson et al. experiments trained large groups of worms in a single rectangular box as a convenience to acquire hundreds of worms for RNA purification, we elected to train our worms in parallel lanes. We did this because isolated worms are more easy to score behaviorally and because James McConnell’s experiments on which Jacobson’s are based primarily used a single-worm narrow lane apparatus with success [64]. Our parallel lane trough allowed us to train multiple worms simultaneously without hampering the visibility of individual worms.

Because our preliminary attempts to replicate the historic protocols showed low and unchanging conditioned response rates to light, we attempted to train a subset of worms that were already reactive to light prior to any training. To do this, we presented a subset of worms with two blocks of 20 light-only trials following the same timing characteristics as described above. Responses to light (turns or contractions) were scored for each trial. The twelve highest-responding worms were split into forward conditioning and pseudoconditioning groups and called “high responders”; the next highest-responding twelve worms were split into the “low responders” groups. Figure S2B shows a box-and-whisker plot of these two groups’ pre-training responses to light.

### 3.4 Contemporary classical conditioning paradigm

Because a group at the University of Tasmania had recently reported success in conditioning planarians using different temporal parameters than our experiments, we attempted to reproduce their procedures [26]. In these experiments, adapted from fear conditioning in rodents, planarians were starved for seven days and subjected to six trials per day for four days, with trials split evenly across one morning and one afternoon block starting at approximately 10:00 and 14:00 respectively. Each trial consisted of 30 seconds of blue light exposure that coterminated with 3 seconds of shock, followed by 30 seconds of rest. The control condition consisted of an approximation of pseudoconditioning, with light-only and shock-only trials randomly dispersed throughout blocks, with at least one of each trial type occurring in each block. The day before training trials began, worms were permitted 10 minutes to explore the trough before being returned to their home dishes. On any training day, worms were given ten minutes to acclimate to the trough before trials began. Other experimental details remained the same as the historic experiments. Data from these experiments were assessed by blinded manual scoring and a computer vision pipeline, described below.

We note two methodological departures from the procedures described by James and colleagues. First, their worms appear to have been trained and tested in circular dishes, whereas ours were trained in linear lanes (as in McConnell’s work [64]) because this enabled us to train multiple worms in one session. Second, while the James group notes that their planarians, described as *G. dorotocephala*, were ordered from a biological supply house, the company that sells them informs us that these worms are collected from ponds in Australia (Southern Biological, personal communication, July 11, 2025). Whether these *G. dorotocephala* worms are invaders from North America, a truly endemic Australian variant, or a different species altogether is unknown. The circular training/test environment and different worm strain could also explain the discrepancy between our results and those of the James group, though further work is needed to resolve this.

### 3.5 Variant conditioning paradigms

Because repeated attempts to condition worms with the historic white light CS and an electric shock UCS (White-Shock) failed, we sought to use other sets of stimuli that might be more salient to planarians. There is some debate as to whether classical conditioning is agnostic to the stimuli used. However, it generally seems that stimuli with special relevance to the natural history of the animal being conditioned are more likely to facilitate learning [68, 69]. For this reason, we added ultraviolet (UV) light to the assembly of stimuli in our training paradigms. We selected ultraviolet light as an additional stimulus because although electric shock is certainly aversive to planarians [70], we are unaware of any ethologically relevant situation where North American planarians would need to respond to electrical shocks. Furthermore, the repeated full-body shock of a submerged organism may cause side effects that could affect worms’ behavior in a way that reflects acute injury rather than learned behavior. In contrast, planarians have dedicated sensory machinery to detect and respond to ultraviolet light, which includes a full-body contraction to sudden onset of intense UV [46, 47, 51]. These morphological and behavioral characteristics are easily reconciled with planarians’ benthic, photophobic lifestyle. Thus, the miscellaneous variant paradigms consisted of combinations of UV light, white light, and shock.

We also varied the species and strain of the worms. In total, we intentionally did not cover the exhaustive space of these stimulus and strain combinations; rather, we sought to rapidly test a subsample for any indication of planarian learning and converge on promising candidate paradigms. Because no promising candidates emerged, we did not pursue these paradigms further.

1. **White-UV**: The same white light from the White-Shock paradigm was paired with an ultraviolet pulse instead of shock. Two *G. dorotocephala* worms collected from Berkeley, California were subjected to 3 seconds of white light coterminating with 1 second of UV light. Worms were trained in three blocks of 25 trials (2-minute ITI), with the first block occurring on Day 1 and the next two blocks on Day 2. A ten minute rest period separated neighboring blocks. Contractions to the onset of white light were scored for each trial. An additional six *S. mediterranea* worms were trained identically, save that they only received two blocks. These experiments took place between 9:30 and 14:30.
2. **UV-Shock**: The white light from White-Shock was swapped with a UV light instead and reduced in strength so as not to trigger baseline contractions. Eighteen *G. dorotocephala* worms were subjected to five blocks of 25 trials of three seconds of UV with onesecond of coterminating shock using an ITI of 2 minutes. Blocks one and two occurred on Day 1, three and four on Day 2, and the fifth on Day 3. Worms were allowed 10–20 minutes to acclimate to their trough lanes before experiments began and given a ten minute rest period between blocks. The shock characteristics were the same 10-12 V, 60 Hz square-wave AC described in our historic protocol. Worms were scored for contractions to the onset of the UV light. These experiments occurred between 10:00 and 16:00.

Because the unconditioned response to a sufficiently high UV stimulus is in itself a contraction, ultraviolet light cannot be considered neutral stimulus in the conventional sense of the term. However, as argued by Jacobson [12], even if conditioned and unconditioned stimuli elicit the same response (contraction), learning can still be assessed by the increasing difference (or lack thereof) in response rates between pseudoconditioning controls and forward/classical conditioning worms. In any case, because we observed no increase in responding among worms trained on any of these paradigms, pseudoconditioning controls were not necessary.

### 3.6 Manual scoring and interscorer agreement

Learning in the classical planarian conditioning assay has historically been measured as the number of full-body contractions that occur in response to the onset of the conditioned light stimulus over the course of training, scored by human observers. In line with this quantification method, we had between one and three scorers manually score each video for conditioned responses occurring after the onset of the conditioned stimulus and before the onset of the unconditioned stimulus. For historic experiments, scorers simply watched complete overhead footage of the experiment as it took place. Because the CS scoring period was only two seconds per trial, each scorer’s response was binary (one contraction or no contraction). By comparing Scorer 1 and Scorer 2’s binary data on a trial-by-trial and worm-by-worm basis, we were able to count the proportion of trials in which each scorer agreed positively (contraction and contraction), agreed negatively (no contraction and no contraction), and disagreed (contraction and no contraction, or vice versa). The proportion of agreement trials divided by the total number of trials is the interscorer agreement for this experiment. We additionally computed Cohen’s *κ* as a measure of interscorer agreement corrected for chance agreement, using the standard two-rater formula *κ* = (*P*_0_ − *P_e_*)/(1 − *P_e_*), where *P*_0_ is the observed proportion of agreement and *P_e_* is the proportion of agreement expected by chance given the marginal response rates of each scorer.

Footage from the contemporary protocol experiments was scored differently using a custom-made computer interface called the Blind Scoring Interface (BSI, Fig. S5). This interface enabled minimally trained scorers to count turns and contractions for each trial of the experimental footage blindly. Three scorers counted conditioned responses across the same subset of 203 trials split between forward conditioning (144) and pseudoconditioning groups (59). Scorer 2 scored an additional thirteen videos (216 total), and Scorer 1 scored all footage of the contemporary paradigm with the interface. Video files representing one worm and one trial were given codenames composed of two random adjectives and one random noun; when a user scored a video, only this codename was seen, so no information about experimental group (forward or pseudoconditioning) was available to them, nor was there any indication of how much previous training the worm had undergone. The videos were created such that each trial containing light presentation was segmented into its own 27-second video spanning the onset of the light up to the frame before the onset of the shock stimulus. Timestamps of turns and contractions were recorded from each video using the blind interface by pressing a “Turn” or “Contraction” button on the frame which the behavior occurred. The video title, as well as the sum and list of event timestamps for turns and contractions, were saved to a CSV from the interface. Data for interscorer reliability and agreement measures were obtained by comparing the event timestamps of turn and contraction events between the three scorers.

Because each scorer scored the same set of videos and noted timestamps of turns and contractions, we were able to compare exactly when each turn and contraction took place. This was accomplished by comparing two scorers’ timestamp lists of turns and contractions per video (one video is equivalent to one trial). For each trial, every possible pair of same-type timestamps was tabulated and the difference between those timestamps calculated. Pairs were sorted by increasing difference and pairs with the least difference were considered matches up to a threshold of ten frames (1 second) for turns and fifteen frames (1.5 seconds) for contractions. Any timestamps remaining when all pair combinations had been exhausted were considered orphans. This was done for same-type events, turns with turns and contractions with contractions, until each type had a list of orphan timestamps. Next, each orphan timestamp was compared to orphan timestamps of the opposite type and, if within an event type’s difference threshold, paired with the opposite-type orphan. Thus, for two scorers’ assessment of a given video, we were able to calculate the number of matching timestamps, the number of unmatched timestamps, and which of those unmatched timestamps could be matched with an opposite-type event. We performed this procedure for each video and for each scorer pair and used those measures to calculate precision, recall and F1 scores.

The F1 metric is the harmonic mean of precision and recall. When treating one scorer’s timestamps as reality (Reference) and testing another scorer against it (Test), precision is the proportion of true positives recovered by the Test dataset out of all Test positives. Recall is the proportion of true Test positives relative to the entirety of the Reference positives. In this context, precision is the proportion of a Test Scorer’s events that can be matched to the Reference Scorer. Likewise, recall is the proportion of Reference events that the Test Scorer can match out of all of the Reference’s events. These proportions describe how much the Test Scorer marks the same events as the Reference Scorer while also not marking events the Reference Scorer does not have. The formulae for precision, recall, and F1 scores are given. (TP = true positive, FP = false positives, FN = false negative.)

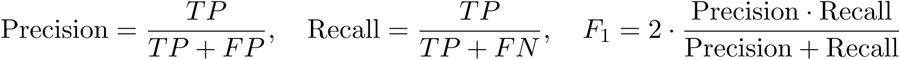

There are two possible variants of F1 score to calculate for scorer pairs. One option is to calculate an F1 score for each video and then average across all video-specific F1 scores to produce a scorer pair’s final F1 score. However, this method risks inflating F1 scores because any video where scorers agree nothing happened (no timestamps given and therefore no possible disagreement) would raise the average F1 score. Instead, we summed the TP, FP, and FN values across all videos, then calculated precision/recall only once to produce a final, more conservative F1 score. The F1 metrics calculated this way are more reflective of agreement about the events of interest.

The measures of unanimous turns and contractions across all three scorers were calculated similarly. For each video, triplets of same-type events were produced and sorted by their range (maximum frame value minus minimum frame value). The range threshold for turns was 10 and 15 for contractions. For each video, some number of unanimous contractions was counted. Because of partial agreement (e.g., Scorer 1 and 2 agreed on a contraction but Scorer 3 disagreed), we used venn diagram notation to calculate the total number of distinct turn and contraction events across all scorers and videos. The sum of all unanimous turns and contractions per video was divided by this number to yield the unanimity rate.

The above methods described agreement about events occurring within the window of the conditioned stimulus, i.e. conditioned responses. To score agreement about contractions induced by shock (the unconditioned stimulus) between Scorers 1 and 2, 198 individual worm trials were sampled from the classical conditioning group of the footage from our attempted contemporary protocol replication. The trials were observed non-blindly for contractions at the onset of the shock stimulus. Blind scoring was deemed unnecessary for shock stimuli because training day or classical/pseudoconditioning status should have no bearing on unconditioned responses barring health issues or habituation, neither of which was observed. The proportion of the number of trials where Scorer 1 and Scorer 2 agreed divided by the number of all scored videos formed the shock agreement metric.

### 3.7 Computer vision pipeline

Because manual scoring was shown to have low interscorer agreement, we sought to develop an alternative method to assess planarian behavior that would be resistant to human bias. The pipeline we developed began by quantifying worm morphology across continuous experimental footage. We used 10 morphological/spatial features (area, perimeter, area-perimeter ratio, circularity, hull area, centroid coordinates, relative concavity area, and two principal components from the 100-dimensional vector of the perimeter shape) to characterize the pose of individual planarians at each frame. To extract these features we tailored the Segment-Anything computer vision pipeline (Meta, Inc) to produce a timeseries of binary masks of the worms from the same overhead experimental footage used to manually score the worms’ behavior. First, each video was spatially cropped such that each video contained a single worm. Then, we labeled two points for the first frame of each video to indicate the worm, and two points to indicate the electrodes (static occlusions in the environment). We used the Segment-Anything pipeline to propagate these points through each video and obtain a timeseries of binary masks. We manually inspected each video, and if the mask extraction quality was suboptimal, split the video before the quality drop and labeled new starting points to propagate through the next video segment. This process was repeated until the entire timeseries of masks was of sufficient quality. Next, we calculated the mask area (sum of all black pixels in the mask), perimeter (using the findContours function from the OpenCV python library), area-perimeter ratio, circularity ((4 ∗ *π* ∗ *A*)/(*P* ∗ *P*) where *A* is the area and *P* is the perimeter), hull area (using convexHull and contourArea from OpenCV python library), centroid coordinates (using moments from OpenCV), and relative concavity area ((*H* − *A*)*/H*) where H is hull area. To complement these features, for each frame we extracted the first two principal components of a 100-dimensional vector of distances from the centroid of the mask to evenly-spaced points on the perimeter of the mask. This analysis was published in [45]. The areas timeseries was used to identify outlier timepoints (*>* 3.5 standard deviations), which were excluded from all feature vectors.

While it is theoretically possible to use the morphological feature information to create a criterion for what counts as a turn or contraction (e.g., a derivative above a certain value in the “Area” timeseries), this approach still relies on human judgment to determine what signature(s) would be suggestive of a conditioned response or merely behavioral noise. The problem inherent to manual scoring is therefore still present. Instead, we took an alternative approach where we quantified how worm behavior differed between conditioned-stimulus (CS-on) and intertrial (CS-off) periods by computing the Kullback–Leibler (KL) divergence between the morphological feature distributions [71]. We first extracted the session-specific windows of time where the conditioned stimulus was on (CS-on) and off (CS-off) and approximated each condition as a multivariate Gaussian with diagonal covariance. We estimated means and unbiased variances of each feature in each condition (ignoring NaNs and adding a small constant for numerical stability), and computed the closed-form 1D Gaussian KL divergence between the CS-on and CS-off Gaussians. We obtained the total divergence KL(CS-on || CS-off) for a given worm during a training block by summing these per-feature KL values. Thus, individual worms’ divergence scores could be plotted to observe any potential change in divergence over the course of the experiment’s four days (eight training blocks).

This computer vision method that assesses learning by the divergence of morphological distributions relies on (1) continuous uninterrupted footage of behaving worms for mask creation, and (2) a sufficient number of frames for each condition (CS-on and CS-off/ITI) to reliably estimate the KL divergence from Gaussian parameters. As described above, the contemporary paradigm contained 30 seconds of CS exposure per trial and an ITI of 30 seconds, which provides ample and continuous footage for analysis. The historic reproduction, conversely, had three-second CS exposure periods with a terminal one-second shock. That provides mere two-second CS-only periods for analysis, which is too little data to rigorously compare to the 30-second ITI. The same issue exists for the variant paradigms. The computer vision method was therefore by necessity restricted to the contemporary paradigm’s footage.

### 3.8 Statistical tests

#### Historic replication

To test for acquisition of the conditioned response over training days, we fit a linear slope to each worm’s daily response rate and tested whether the mean slope differed from zero (one-sample *t* -test, two-sided). We confirmed this with Page’s L test for monotone increasing trends across ordered time points. Group differences (conditioning vs. pseudoconditioning) were assessed with a one-way ANOVA. All analyses used Python (SciPy, statsmodels).

#### Contemporary replication

To test for acquisition of the conditioned response from the hand-scored data, we computed a linear slope of each worm’s daily response count across training days and tested whether the mean slope was significantly greater than zero using a one-sample *t* -test (one-sided, *α* = 0.05). We supplemented this with Page’s L test, and fit LMMs with Day (continuous), Condition (conditioning vs. pseudoconditioning), and their interaction as fixed effects and individual subject as a random intercept, estimated via maximum likelihood. The Day × Condition interaction served as the primary test of the conditioning hypothesis, assessing whether the rate of behavioral change over training days differed between groups. LMMs were fit separately for turns and contractions. To compare each subpopulation’s mean per-worm slope to the historic reference, we used Welch’s *t* -tests with denominator degrees of freedom adjusted to propagate the standard error of Jacobson’s published regression slope.

#### Contemporary protocol scoring with the computer vision pipeline

To test for changes in KL divergence values between conditioning and pseudoconditioning groups over training, we log-transformed the KL divergence values to improve normality and analyzed with a LMM predicting log KL from Day, Condition, and their interaction, with random intercepts for individual subjects. The Day × Condition interaction term served as the primary test of differential learning trajectories; the main effect of Condition was additionally examined to assess overall group differences in KL divergence level. Per-subject slopes were also compared between conditions using Welch’s *t* -test.

#### Miscellaneous conditioning paradigms

For each paradigm (UV-shock and white light–UV), we computed a linear slope of each worm’s contraction count across training blocks and compared each population’s mean per-worm slope to the historic Jacobson reference of +1.40 responses/block using two-sided Welch’s *t* -tests with denominator degrees of freedom adjusted to propagate the standard error of Jacobson’s published regression slope, and computed one-sample Cohen’s *d* against the fixed Jacobson reference value. Worms were pooled across cohorts within each paradigm; slopes from cohorts contributing different numbers of blocks were weighted equally at the worm level.

#### Individual sensitivity

To test if different sensitivity to the conditioned stimulus affected acquisition of the conditioned response, we computed each worm’s linear slope across conditioning blocks and tested whether the mean slope differed from zero (one-sample *t* -test, two-sided), separately for the conditioning and pseudoconditioning groups. To confirm that the high and low responder groups were meaningfully separated at baseline, we compared their pre-training CS response rates using an independent-samples *t* -test and computed Cohen’s *d* as a measure of effect size. Slopes were compared between high and low responders within each condition using Welch’s *t* -test with Cohen’s *d*. To assess whether response rates changed across the light-only assessment sessions, we compared per-worm Day 1 and Day 2 means using a paired *t* -test (two-sided) with Cohen’s *d_z_*. To distinguish a gradual within-session trend from a between-session shift, we additionally fit per-worm linear slopes across individual trials using a one-sample *t* -test and computed Spearman’s rank correlation between trial number and response rate.

## Supporting information

**Fig S1.**
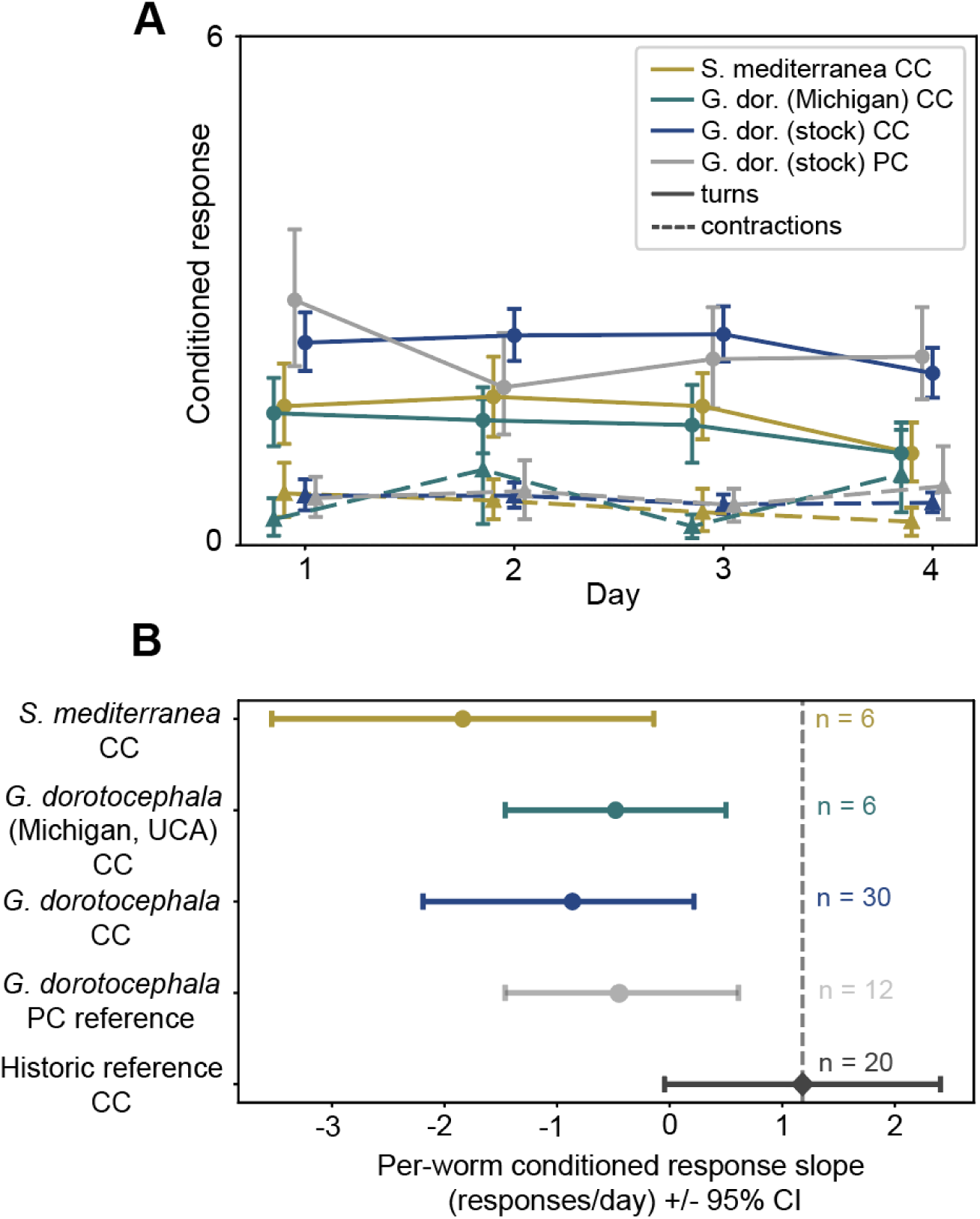
Conditioned-response acquisition from contemporary conditioning protocol, by species and group. *A*. Mean per-trial conditioned response score per worm per day, for turns (solid) and contractions (dashed), broken down by species and strain (data from Fig. 2B). Error bars indicate 95% bootstrap CIs on the daily mean. One PC reference group (stock *G. dorotocephala*) shown for context. *B*. Forest plot of population-mean per-worm OLS slopes (± 95% CI) for three CC populations and two references. Slopes pool turns and contractions. Dark-gray diamond: Jacobson’s historic CC reference slope of +1.40 ± 0.40 SE responses/day, derived by OLS on five published block means.

**Fig S2.**
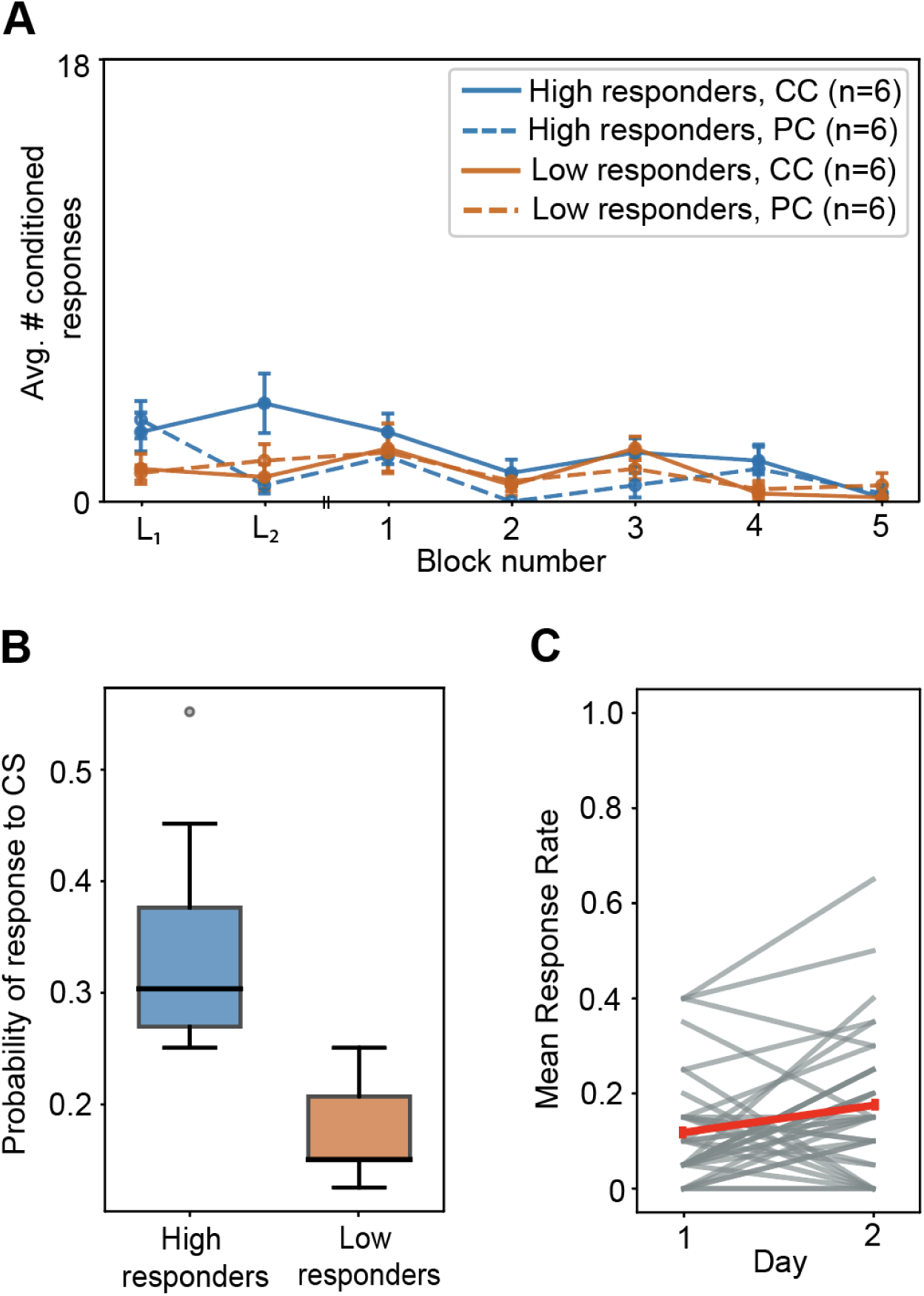
Pre-conditioning baseline characterization and per-block conditioned response acquisition for historical protocol cohort. *A*. Mean conditioned responses per block, by baseline-responsiveness stratum and condition (*n* = 6 per line; error bars: 95% bootstrap CI). Note that the CC and PC groups did not differ in overall response level, and slopes did not differ between strata within either condition. *B*. Per-worm mean CS response rate across two baseline days of exposure to the CS only (*n* = 12 per stratum). Box edges: 25th–75th percentiles; line: median; whiskers: 1.5 × IQR. High and low strata were significantly separated at baseline (*t*(22) = 5.58*, p < .*001, Cohen’s *d* ≈ 2.28). *C*. Per-worm mean response rate across the two-day light habituation test (*N* = 36; grey lines: individuals; red line: grand mean ± SEM). No within-session trend was detected; response rate increased between sessions, contrary to habituation (+1.5 × 10*^−^*^4^ ± 9.3 × 10*^−^*^4^ resp./trial, *t*(35) = +0.16*, p* = .873; Spearman *ρ* = +0.002*, p* = .95).

**Fig S3.**
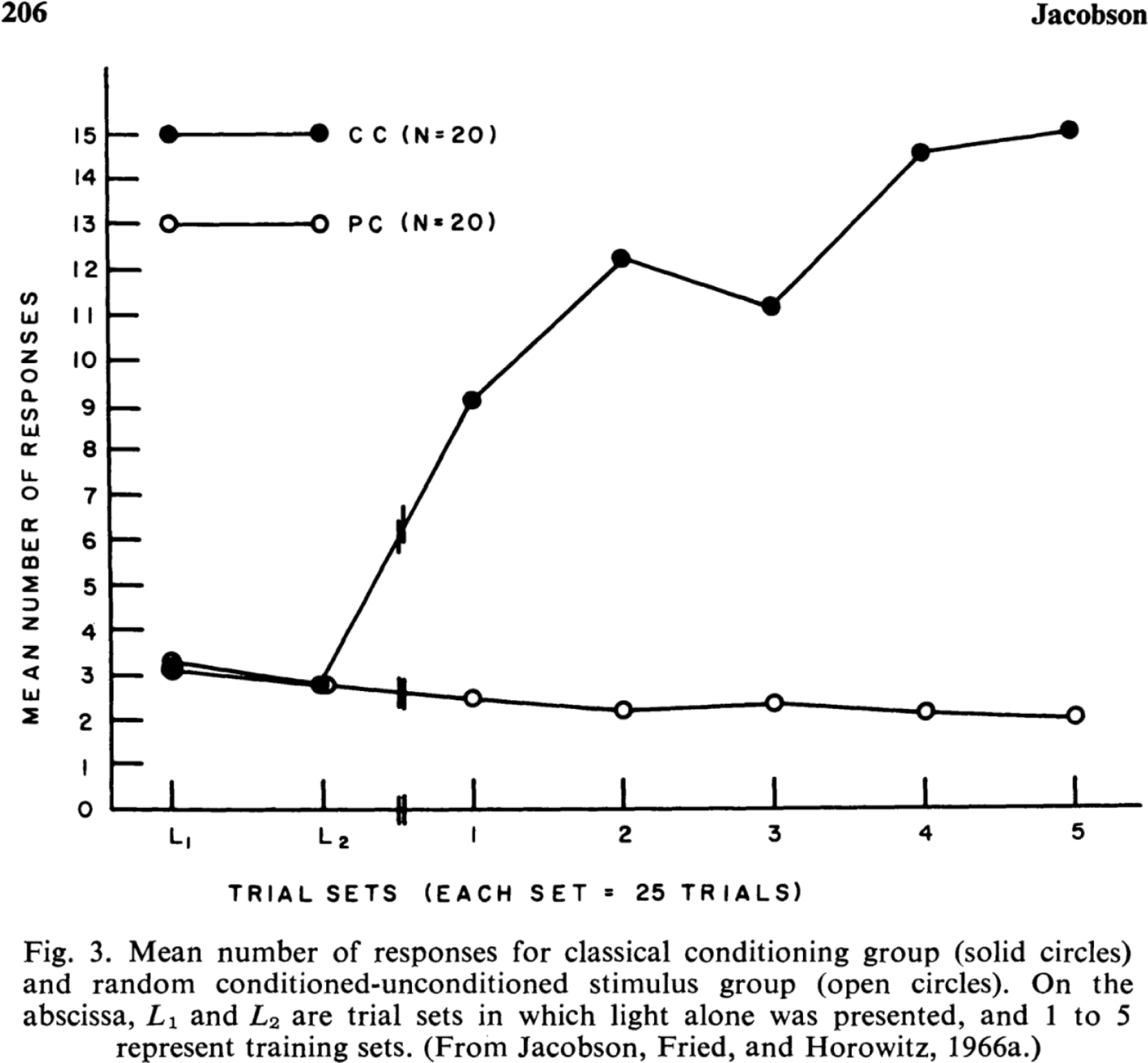
Reprinted original plot from Jacobson et al. 1967.

**Fig S4.**
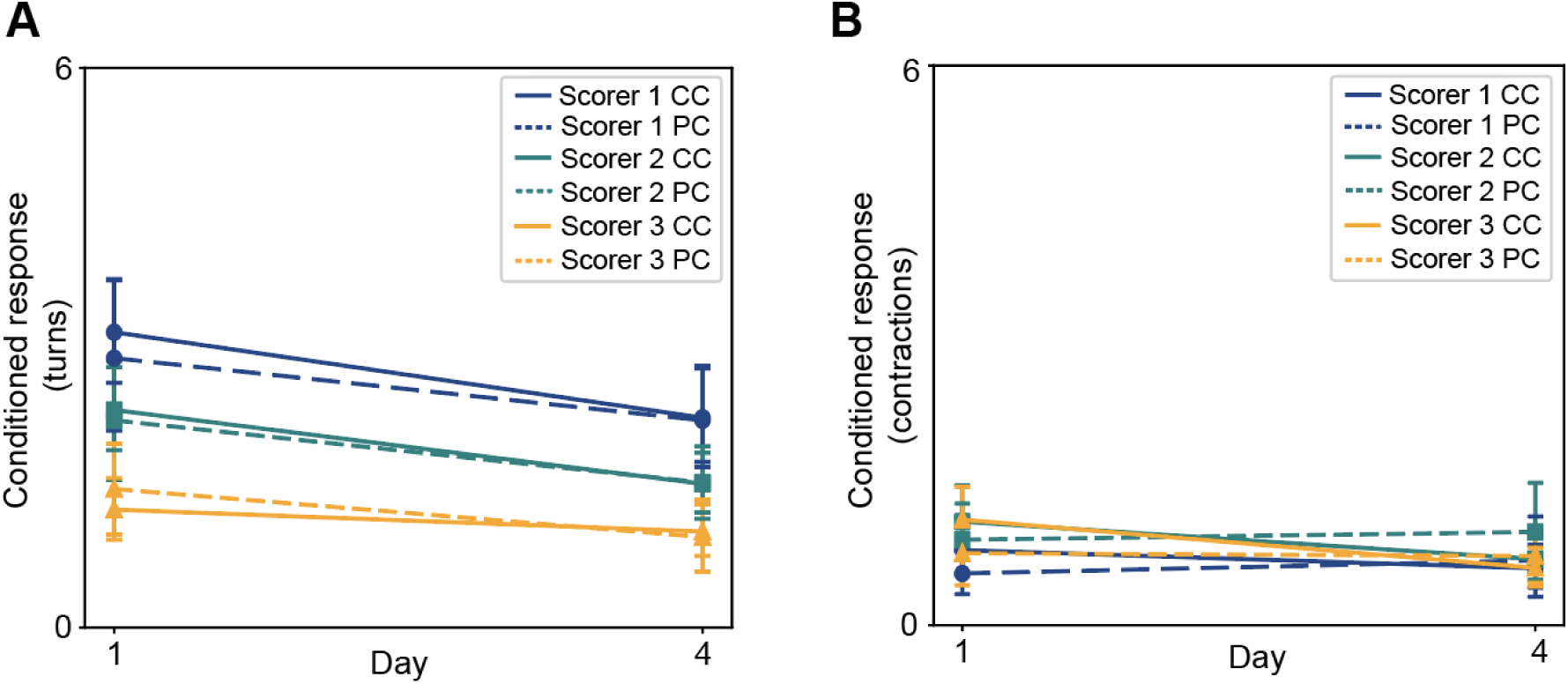
Three independent blind scorers all fail to detect an increase in conditioned responses during the contemporary conditioning protocol. *A*. Across-worm mean turn count on Day 1 and Day 4 for each scorer (CC: solid lines, 72 worm-sessions per scorer × day, with Scorer 3 at n = 65-68 due to missing sessions); PC: dashed lines, n = 36 worm-sessions per scorer × day, with Scorer 3 at n = 35). Error bars indicate 95% bootstrap CIs. No scorer shows an increase from Day 1 to Day 4 in either group. *B*. Same as A, for contraction responses.

**Fig S5.**
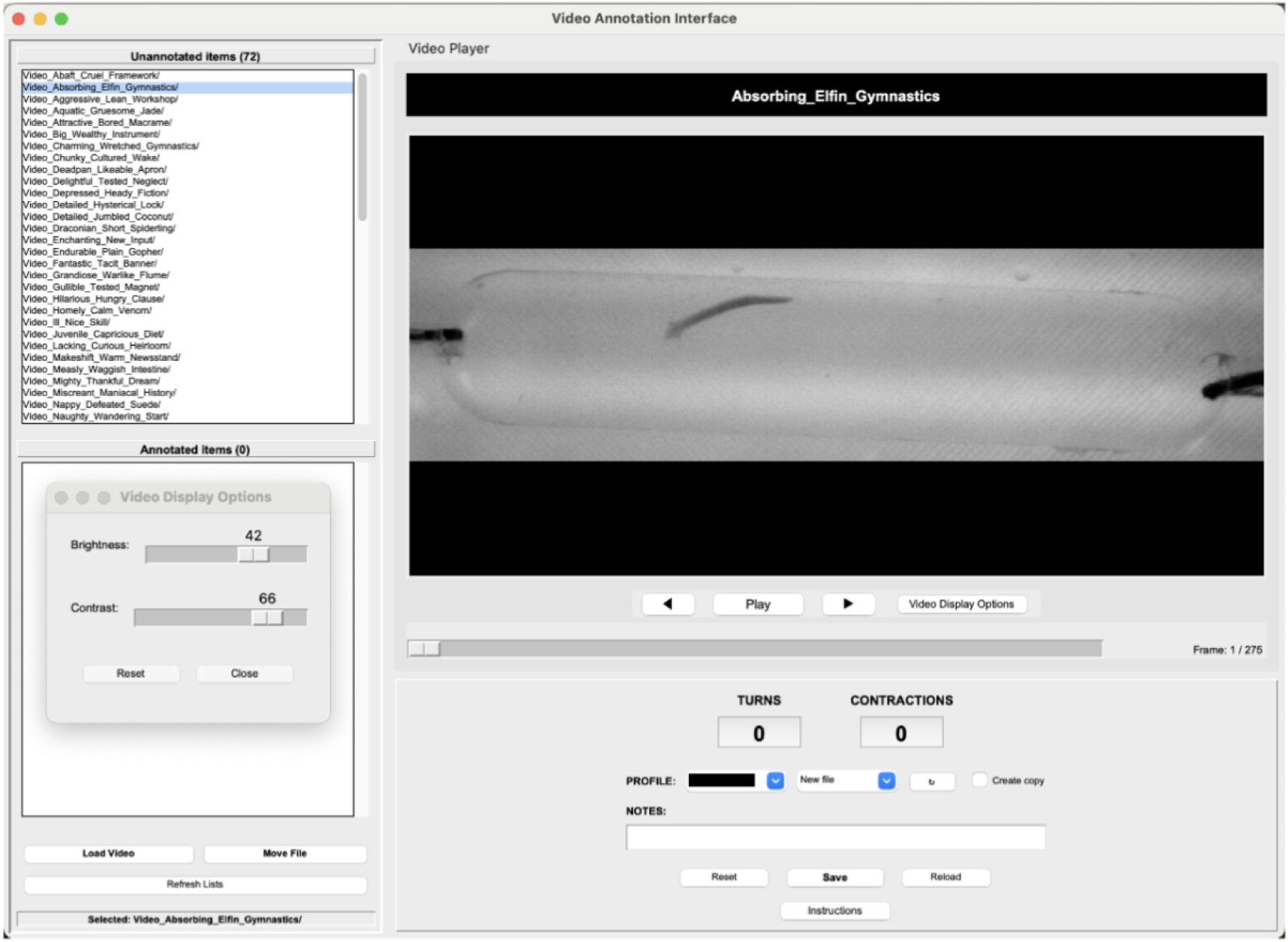
Blind Scoring Interface. Graphical user interface used to score worm behavior from the contemporary protocol. Each trial was cropped to a 27 s video spanning CS onset to just before UCS onset and assigned a random codename. Scorers marked turn and contraction events without knowledge of the worm’s experimental group or training history.

## Notes

https://github.com/madeleinesnyder/PlanariaPipe/tree/main

https://doi.org/10.7910/DVN/NSTJGZ

### Competing Interest Statement

The authors have declared no competing interest.

### Summary of Updates

The revision has been revised to strengthen methodological clarity, qualify statistical claims according to sample size, and bring clarity to the aims to of the paper.

### Funding

- Schmidt Sciences

