## Supplementary information - revised for "Failure to classically condition planarian flatworms"

### Supporting information

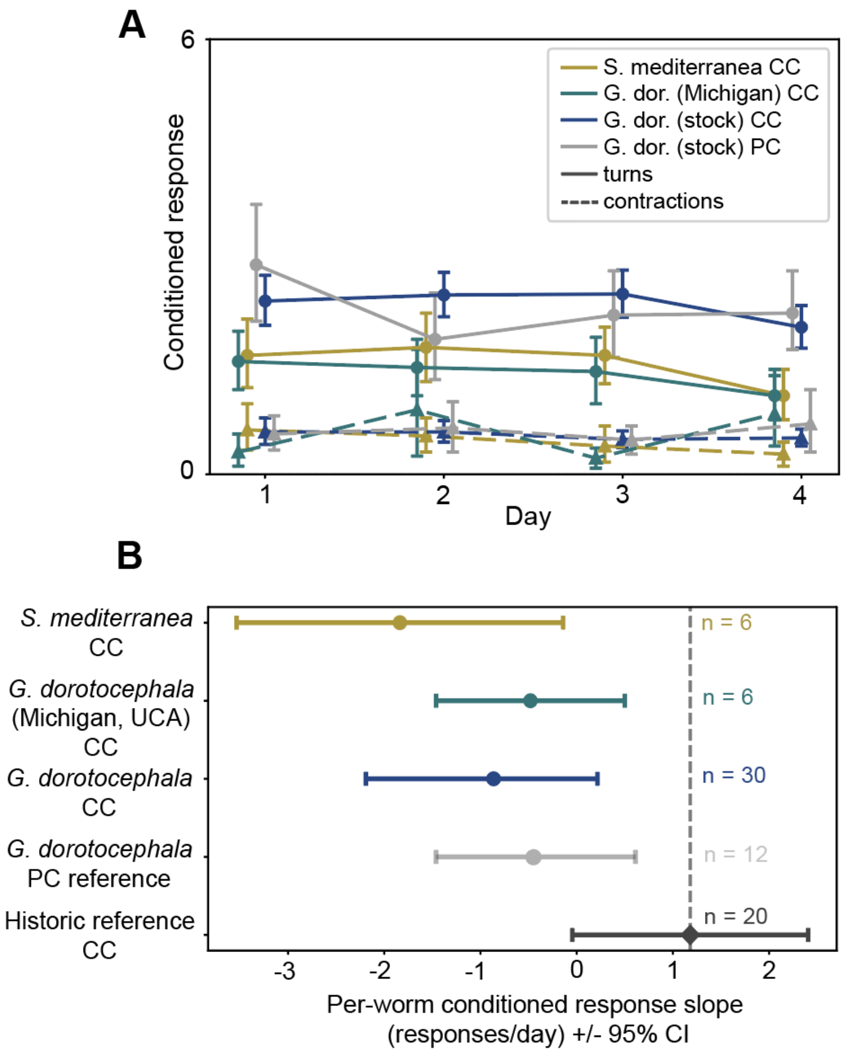


**Fig S1. Conditioned-response acquisition from contemporary conditioning protocol, by species and group.** *A*. Mean per-trial conditioned response score per worm per day, for turns (solid) and contractions (dashed), broken down by species and strain (data from Fig. 2B). Error bars indicate 95% bootstrap CIs on the daily mean. One PC reference group (stock *G. dorotocephala*) shown for context. *B*. Forest plot of population-mean per-worm OLS slopes (± 95% CI) for three CC populations and two references. Slopes pool turns and contractions. Dark-gray diamond: Jacobson's historic CC reference slope of $+1.40 \pm 0.40$ SE responses/day, derived by OLS on five published block means.

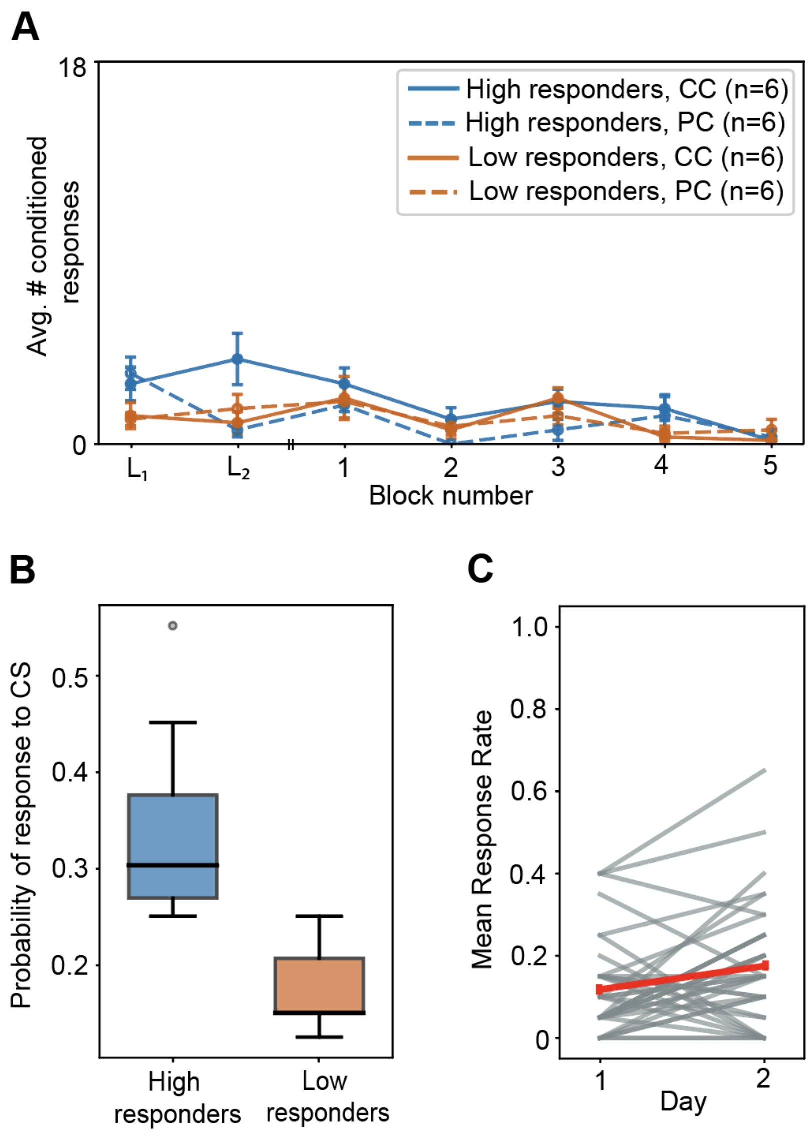


**Fig S2. Pre-conditioning baseline characterization and per-block conditioned response acquisition for historical protocol cohort.** *A.* Mean conditioned responses per block, by baseline-responsiveness stratum and condition ($n = 6$ per line; error bars: 95% bootstrap CI). Note that the CC and PC groups did not differ in overall response level, and slopes did not differ between strata within either condition. *B.* Per-worm mean CS response rate across two baseline days of exposure to the CS only ($n = 12$ per stratum). Box edges: 25th–75th percentiles; line: median; whiskers: 1.5 × IQR. High and low strata were significantly separated at baseline ($t(22) = 5.58, p < .001$, Cohen's $d \approx 2.28$). *C.* Per-worm mean response rate across the two-day light habituation test ($N = 36$; grey lines: individuals; red line: grand mean ± SEM). No within-session trend was detected; response rate increased between sessions, contrary to habituation ($+1.5 \times 10^{-4} \pm 9.3 \times 10^{-4}$ resp./trial, $t(35) = +0.16, p = .873$; Spearman $\rho = +0.002, p = .95$).

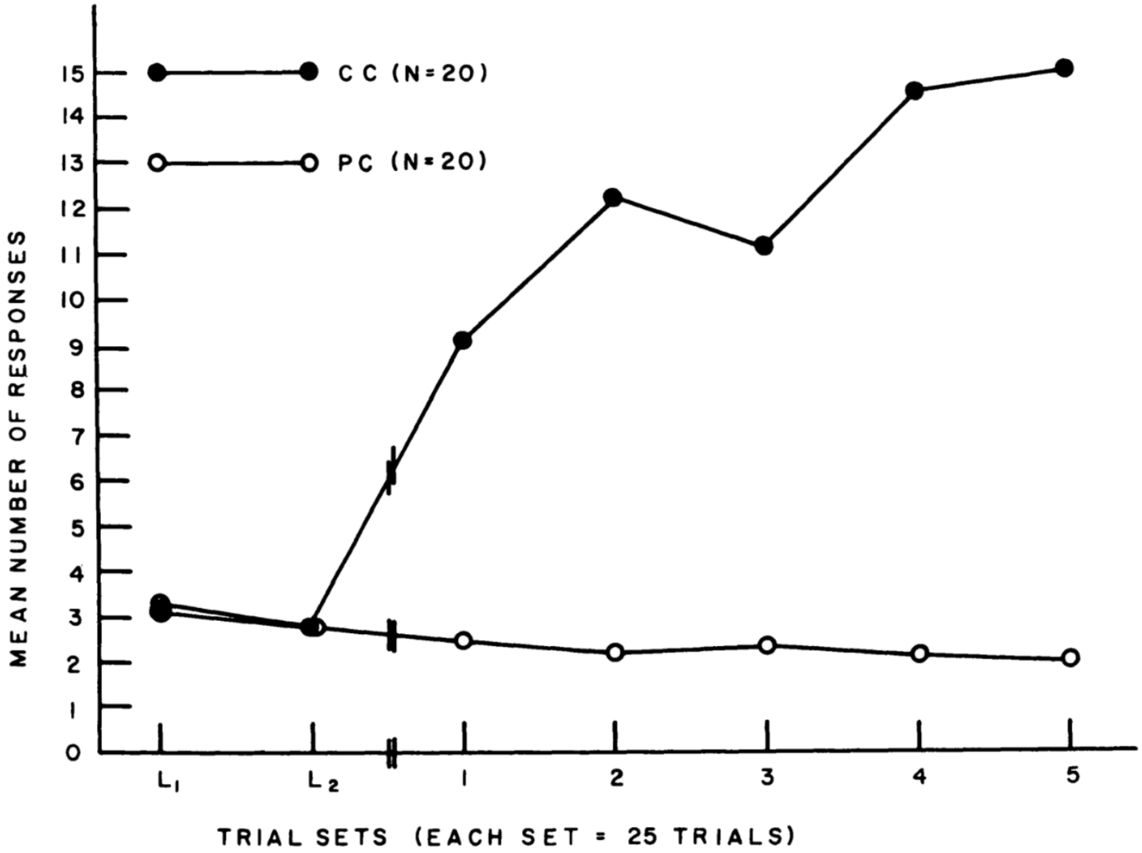


Fig. 3. Mean number of responses for classical conditioning group (solid circles) and random conditioned-unconditioned stimulus group (open circles). On the abscissa, $L_1$ and $L_2$ are trial sets in which light alone was presented, and 1 to 5 represent training sets. (From Jacobson, Fried, and Horowitz, 1966a.)

**Fig S3. Reprinted original plot from Jacobson et al. 1967.**

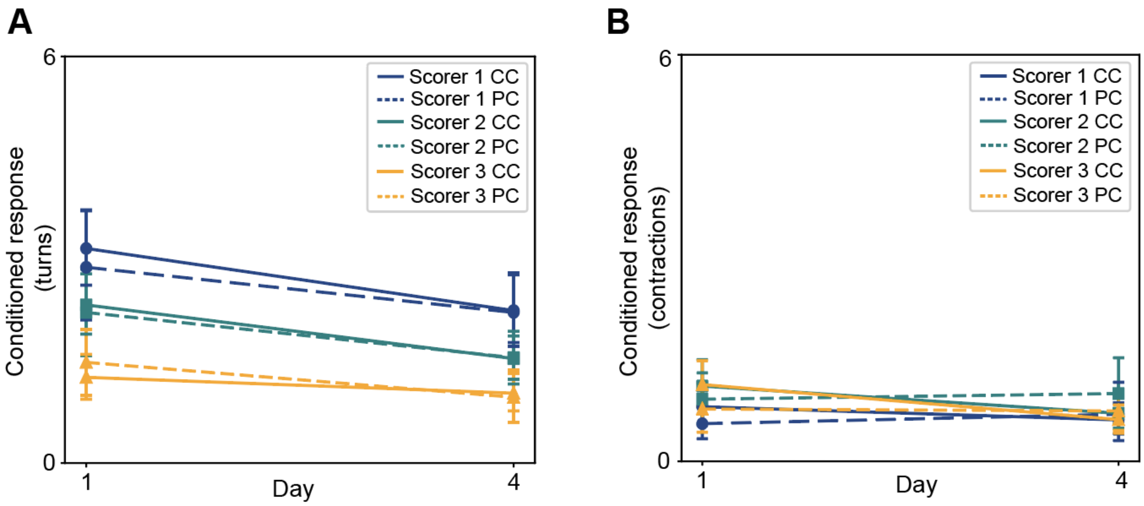


**Fig S4. Three independent blind scorers all fail to detect an increase in conditioned responses during the contemporary conditioning protocol.** *A*. Across-worm mean turn count on Day 1 and Day 4 for each scorer (CC: solid lines, 72 worm-sessions per scorer × day, with Scorer 3 at n = 65-68 due to missing sessions); PC: dashed lines, n = 36 worm-sessions per scorer × day, with Scorer 3 at n = 35). Error bars indicate 95% bootstrap CIs. No scorer shows an increase from Day 1 to Day 4 in either group. *B*. Same as A, for contraction responses.

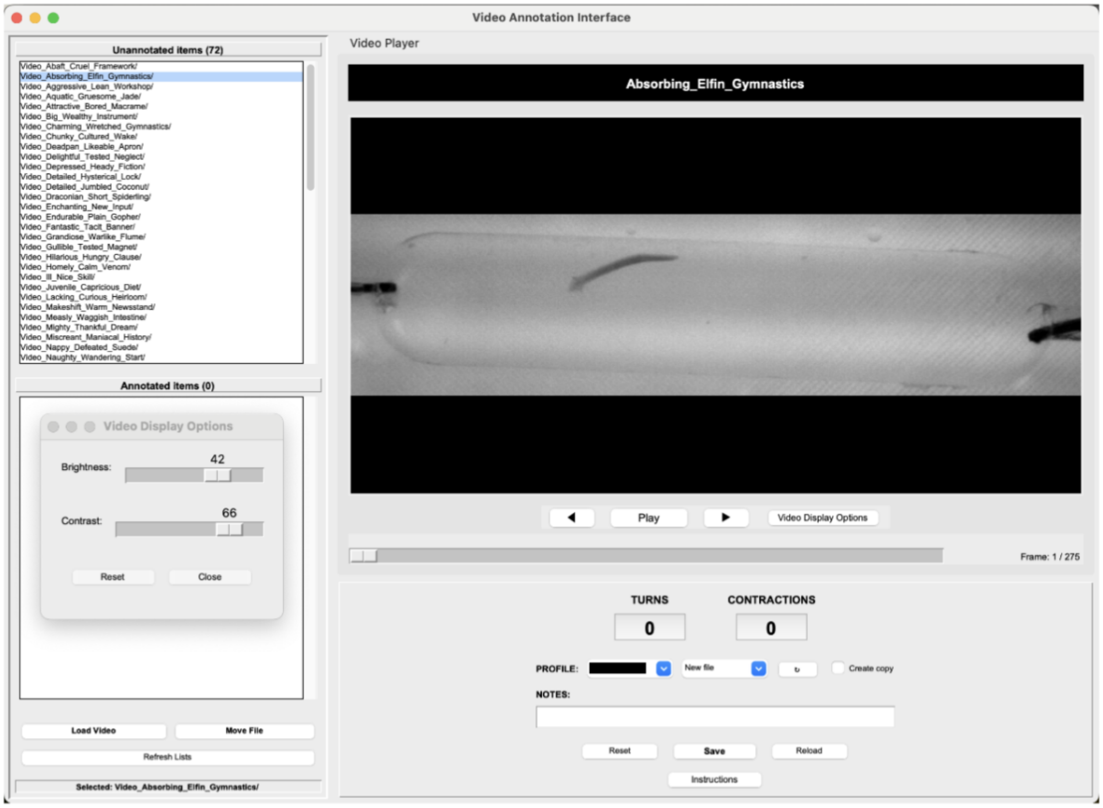


**Fig S5. Blind Scoring Interface.** Graphical user interface used to score worm behavior from the contemporary protocol. Each trial was cropped to a 27 s video spanning CS onset to just before UCS onset and assigned a random codename. Scorers marked turn and contraction events without knowledge of the worm's experimental group or training history.
